# TurboID-based proteomic profiling reveals proxitome of the IRT1 metal transporter and new insight into metal uptake regulation in plants

**DOI:** 10.64898/2026.03.16.712057

**Authors:** Liza Pellegrin, Steven Fanara, Bertrand Fabre, Carole Pichereaux, Valérie Cotelle, Grégory Vert, Julie Neveu

**Affiliations:** Laboratoire de Recherche en Sciences Végétales (LRSV), CNRS, Université de Toulouse, CNRS, Toulouse-INP, Auzeville-Tolosane, France; Institut de Pharmacologie et de Biologie Structurale (IPBS), CNRS, Université de Toulouse, Toulouse, France; Fédération de Recherche Agrobiosciences, Interactions et Biodiversité (FRAIB), CNRS, Université de Toulouse, Auzeville-Tolosane, France; Infrastructure Nationale de Protéomique, ProFI, UAR 2048, Toulouse, FR, France

**Keywords:** Arabidopsis, Iron, Metal, IRT1, endocytosis, TurboID, ubiquitin, transporter

## Abstract

IRT1 is the major root iron transporter responsible for broad spectrum metal absorption in Arabidopsis root epidermal cells. Non-iron metal substrates of IRT1 were recently shown to regulate IRT1 cell surface levels by endocytosis and vacuolar degradation. TurboID-based proximity labeling was recently developed to detect protein:protein interactions and thus shed light on the intricate regulation of proteins in living cells. Although TurboID-based proximity labeling technology has been successfully established in mammals, its application in plant systems remains limited and inexistent for highly hydrophobic multispann transmembrane proteins. Here, we established TurboID for proximity labeling of IRT1 and identified 494 IRT1-specific proximal proteins, including the previously reported FYVE1 IRT1 interacting protein. To showcase the biological relevance of identified IRT1 proximal proteins, we characterized further the NHX5 Na^+^(K^+^)/H^+^ antiporter and the RGLG2 E3 ubiquitin ligase. We validated both IRT1 proximal proteins as IRT1 partners using several orthogonal assays. We also highlight the contribution of NHX-type antiporters and RGLG-type E3 ligases in plants responses to non-iron metal nutrition and IRT1 endocytosis. Overall, our work showcases the power of TurboID to identify new interacting proteins for plant transporters, expanding the application of this technology to proteins notoriously difficult to work with.

## Introduction

Iron is essential for plant growth and development, playing fundamental roles in many cellular processes including photosynthetic and respiratory electron transfer reactions. Conversely, overload of iron is also toxic, leading to oxidative stress. Soil iron bioavailability to plants is often limited, such as in calcareous soils where iron is present under the form of insoluble complexes (Briat et al., 1995). To maintain iron homeostasis, plants must tightly regulate iron absorption from the soil. In non-graminaceous plants, including the model plant *Arabidopsis thaliana*, iron absorption by root epidermal cells is achieved through the so-called strategy I that requires three successive steps. First, soil ferric chelates are solubilized by local rhizosphere acidification, notably via the release of protons by the proton pump H^+^-ATPase2 (AHA2). Solubilized Fe^3+^ ions are then reduced to Fe^2+^ by the Ferric Reduction Oxydase2 (FRO2) reductase and finally transported into the cell by the iron transporter Iron Regulated Transporter1 (IRT1) (Palmer and Guerinot, 2009; Thomine and Vert, 2013; Jeong et al., 2017). Apart from the predominant role of IRT1, FRO2 and AHA2, another membrane protein called ATP-Binding Cassette G37 (ABCG37/PDR9) was demonstrated to be involved in Arabidopsis iron acquisition by exporting coumarins in the rhizosphere under iron deficiency (Fourcroy et al., 2014). These excreted phenolic compounds chelate Fe^3+^ and facilitate iron availability for reduction by FRO2 (Fourcroy et al., 2016).

IRT1 is a major player in the regulation of plant iron homeostasis, as illustrated by the severe chlorosis and the lethality of *irt1-1* knock-out mutant (Vert et al., 2002). As such, IRT1 is at the center of complex transcriptional regulatory networks aiming at integrating plant iron nutrition with several endogenous and exogenous cues. In Arabidopsis, the expression of iron uptake genes is activated under iron-limited conditions through an intricate cascade of basic Helix-Loop-Helix (bHLH) transcription factors and E3 ubiquitin ligases, culminating in the strong accumulation of *IRT1* and *FRO2* transcripts in root epidermal cells. Various transcription factors, such as ERFs and WRKYs, or phytohormones like salicylic acid, gibberellin, jasmonic acid, auxin, and ethylene, have also been shown to impact on the transcriptional regulation of *IRT1* to adjust plant iron nutrition to growth conditions (Riaz and Guerinot, 2021; Zhang et al., 2023; Mahawar et al., 2023).

Recently, much progress has been made in our understanding of *IRT1* post-translational regulation and function at the cell surface. IRT1 is found at the plasma membrane, where it associates with FRO2 and AHA2 to mediate iron uptake, in the *trans* Golgi Network/early endosomes (TGN/EE), and is targeted to the vacuole for degradation (Barberon et al., 2011; Martín-Barranco et al., 2020). IRT1 was shown to undergo constitutive endocytosis, using AP2 adapters and recognition of canonical Tyr-based motifs within its regulatory loop, and cycling back to the cell surface (Barberon et al., 2011; Spielmann et al., 2025). IRT1 is a broad-spectrum transporter that takes up zinc, manganese, cobalt and cadmium (Zn, Mn, Co, or Cd hereafter called non-iron metal substrates), in addition to iron (Rogers et al., 2000; Vert et al., 2001; Vert et al., 2002), and IRT1 endocytosis is triggered by excess of such non-iron metal substrates (Barberon et al., 2014; Dubeaux et al., 2018; Spielmann et al., 2022). IRT1 directly binds non-iron metals using histidine-rich stretch located in IRT1 large cytosolic loop with the help of a nearby aspartic acid residue (Dubeaux et al., 2018; Spielmann et al., 2022; Cointry et al., 2025). Non-iron metal binding to IRT1 allows the recruitment of Calcineurin B-like (CBL)-interacting serine/threonine-protein kinase 23 (CIPK23) and subsequent phosphorylation of IRT1 (Dubeaux et al., 2018). This in turn disrupts the association of IRT1 with AHA2 and FRO2 (Martín-Barranco et al., 2020), and allows lysine-63 polyubiquitination of IRT1 by IDF1 and subsequent targeting to the vacuole (Dubeaux et al., 2018). Such elegant regulatory mechanism protects plants from overaccumulating readily available Zn^2+^, Mn^2+^, Co^2+^ and Cd^2+^ ions that do not need prior reduction by FRO2 to be transported. IRT1 is found in the outer plasma membrane domain facing the rhizosphere, which is necessary for proper radial transport of metals in the root (Barberon et al., 2014). IRT1 physically interacts with both the late-endosome-localized ESCRT-associated FYVE1 (Barberon et al., 2014; Gao et al., 2014) and AP2 at the cell surface (Spielmann et al., 2025) to maintain its lateral polarity, pointing to a role of recycling and internalization in this process. IRT1 recycling from endosomes to the plasma membrane was also shown to require Sorting Nexin 1 (SNX1) (Ivanov et al., 2014). Finally, the peripheral membrane protein ENHANCED BENDING1 (EHB1) was demonstrated to interact with IRT1 in a calcium-dependent manner and was proposed to act as an inhibitor of IRT1-mediated iron transport (Khan et al., 2019).

IRT1 was established as a model to study the post-translational regulation of plasma membrane proteins and its functional relevance through the identification of IRT1 partners. To date, identifying partners of membrane proteins including highly hydrophobic multispan transmembrane transporters and channels remains challenging. Affinity purification coupled to mass spectrometry (AP-MS) fails to capture weak and transient interactors and requires membrane protein solubilization with mild detergents not to interfere with affinity purification, thus yielding many false positives. Yeast two hybrid and split-ubiquitin are tedious, require the use of soluble fragments (Y2H), are also prone to false positives, and suffer from the use the yeast heterologous system. The recent development of proximity labeling, such as BioID, to capture protein-protein interactions *in planta* overcomes many caveats from traditional approaches, especially for membrane proteins. BioID uses a protein of interest (POI) fused to a promiscuous biotin ligase to catalyze biotinylation of amine groups on lysine residues from nearby proteins, usually within a radius of about 10 nm. The biotinylated proximal proteomes (i.e. proxitomes) of the POI can be enriched using streptavidin-based purification under stringent conditions followed by mass spectrometry-based proteome identification.

To shed light on new actors contributing to iron uptake and new regulatory proteins, we identified the proxitome of the IRT1 transporter. To this purpose, we pioneered BioID proximity labeling approaches for an integral multispanning hydrophobic plant membrane protein. We uncovered the IRT1 proxitome under different metal regimes, and validated two candidate proximal proteins as new IRT1-interacting proteins and players in plant metal homeostasis. The BioID-based proximity labeling strategy detailed in this study sets the stage to identify partners for other transporters and channels to better understand their role and regulation.

## Results

### Implementing BioID for the IRT1 metal transporter

To carry out proximity labelling approaches using IRT1 through BioID, we first thought of generating a transgenic line expressing a functional fusion between IRT1 and the biotin ligase TurboID. TurboID has indeed emerged as the most efficient labelling enzyme in various plant systems (Branon et al., 2018; Mair et al., 2019). IRT1 is a highly hydrophobic multipass transmembrane protein with extracellular N- (after cleavage of the predicted signal peptide) and C-termini, and the only functional fusion of IRT1 with a tag of significant size reported to date harbors the tag inserted in the first extracellular domain of IRT1 (Dubeaux et al., 2018). Since our goal was to use TurboID to identify intracellular partners, we screened for functional IRT1-TurboID fusions in which the biotin ligase is located in the cytosolic regions of IRT1, while maintaining IRT1 function. To this purpose, we generated several IRT1-TurboID fusions, where TurboID is inserted in different cytosolic loops, under the control of the *IRT1* promoter in the *irt1* mutant background. Several independent monoinsertional homozygous lines for the different fusions were isolated. Among all the lines generated, the one carrying TurboID in the first cytosolic loop of IRT1 between transmembrane domain (TM) 1 and 2 (between residues S73 et R74 ; Figure 1A) complemented the severe chlorosis of *irt1*, thus attesting that such fusion is still functional for iron transport (Figure 1B). In parallel, we confirmed that the generic plasma membrane protein Lti6b failed to interact with IRT1 as observed by the lack of coimmunoprecipitation of endogenous IRT1 with Lti6b-GFP (Figure 1C). We therefore generated a TurboID negative control line expressing the Lti6b fused to TurboID under the control of the *IRT1* promoter in the wild-type background.

**Figure 1.**
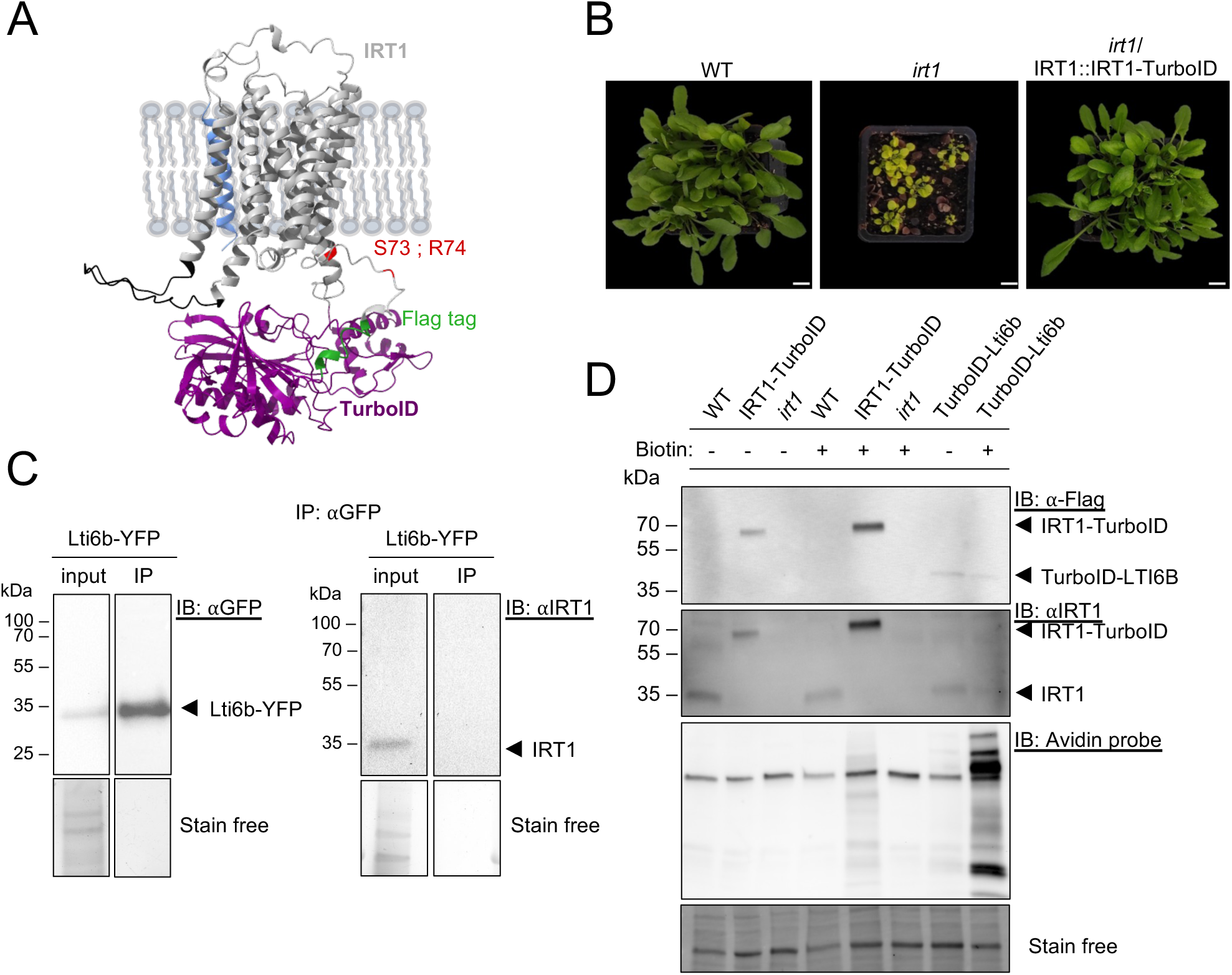
Strategy for establishing TurboID approach for the IRT1 metal transporter. (A) Alphafold-based model prediction of the IRT1-TurboID fusion protein. IRT1, grey (with putative signal peptide in blue) ; TurboID, purple ; Flag tag, green. The TurboID insertion site (between S73 and R74) is shown in red. (B) Phenotype of 4-week-old wild-type (WT, Col), *irt1* crispr mutant, and complemented *irt1*/IRT1::IRT1-TurboID plants. (C) Immunoprecipitation of Lti6b-YFP. Immunoprecipitation was performed using anti-GFP antibodies on solubilized root protein extracts from 35S::Lti6b-YFP plants and subjected to immunoblotting with anti-GFP (left) or anti-IRT1 antibodies (right). Plants expressing Lti6b-YFP were grown for 14 days on iron-deficient conditions prior to protein extraction. Inputs and immunoprecipitated fractions are shown. IB, immunoblotting; IP, immunoprecipitation. The stain free signal is used as loading control. (D) Characterization of IRT1-TurboID and TurboID-Lti6b lines. Western blot analyses monitoring IRT1, IRT1-TurboID/TurboID-Lti6b and biotinylated protein accumulation in wild-type (WT), *irt1*, *irt1*/IRT1::IRT1-TurboID and IRT1::TurboID-Lti6b using anti-IRT1 antibodies, anti-FLAG antibodies, or avidin probe, respectively. Total proteins were extracted from roots of 14-day-old plants grown in half-strength MS media, treated with 100 µM Ferrozine for 24 hours, and subjected to with mock or 200 µM biotin in liquid medium for 4 hours. The stain free signal is used as loading control. (E) Influence of biotin addition of IRT1-TurboID ubiquitination profile. Immunoprecipitation of IRT1-TurboID was performed on root protein extracts probed using anti-ubiquitin antibodies (P4D1). Proteins were extracted from 14-day-old plants expressing IRT1::IRT1-TurboID treated with non-iron metals for 2 hours in liquid medium prior to addition of 200 µM biotin in non-iron metal liquid medium. (F) Alt Text : Illustration and data on the generation and characterization of IRT1 transgenic lines for TurboID.

To characterize these lines and ascertain that the IRT1-TurboID cassette is expressed at levels close to endogenous IRT1, we first performed western blot analyses using total proteins extracted from roots. Since the IRT1-TurboID and Lti6b-TurboID cassettes carry a Flag tag, protein detection was performed using anti-Flag antibodies. The IRT1-TurboID protein nicely accumulated while Lti6b-TurboID accumulated only to low levels (Figure 1D, upper panel). To compare IRT1-TurboID levels to endogenous IRT1 levels from wild-type plants, protein extracts were probed with anti-IRT1 antibodies. This proved that IRT1-TurboID accumulated to similar levels than endogenous IRT1 protein (Figure 1D, middle panel), pointing to the relevance of such line to probe for IRT1 proxitome using BioID.

The fact that the biotin ligase is inserted between 2 TM may potentially affect its ligase activity. We therefore tested the labelling efficiency of TurboID in IRT1-TurboID and Lti6b-TurboID lines. Plants were exposed to 50 µM biotin or mock for 2 hours, and total protein biotinylation was detected by immunoblot using avidin probes. Biotin addition greatly increased protein biotinylation in IRT1-TurboID lines compared to both *irt1* and wild-type plants, attesting of the functionality of the TurboID ligase (Figure 1D, bottom panel). Interestingly, Lti6b-TurboID showed strong biotinylation, although the corresponding fusion protein accumulated only at low levels, indicating that harboring TurboID at the protein extremity imposes less physical constraints and yields higher activity (Figure 1D, bottom panel). Considering that biotin was previously reported to coordinate metals, we also ascertained that biotin treatment did not interfere with *IRT1* expression and localization in response to metals (Supplemental Figure 1A).

We then assessed if protein biotinylation impacted on the previously reported ubiquitination and endocytosis of IRT1 triggered by non-iron metal excess (Dubeaux et al., 2018). This is all the more important considering that covalent binding of biotin to protein uses lysine residues that can also be targeted by ubiquitin. IRT1 ubiquitination profiles were therefore determined by immunoprecipitating IRT1-TurboID from plants initially grown in the absence of iron and non-iron metals, and then subjected to simultaneous combinations of non-iron metals and biotin, before detection with anti-ubiquitin antibodies. As expected, low ubiquitination of IRT1-TurboID was detected in absence of non-iron metals (Supplemental Figure 1B). When exposed to non-iron metals, IRT1-TurboID showed increased ubiquitination (Supplemental Figure 1B), as previously observed (Dubeaux et al., 2018). However, simultaneous biotin addition resulted in biotinylation of IRT1-TurboID itself (Supplemental Figure 1B), preventing IRT1 ubiquitination because of the likely competition for target lysine residues. Alternatively, the presence of numerous lysine residues within ubiquitin that are likely biotinylated may also impair ubiquitin recognition by anti-ubiquitin antibodies. To circumvent this caveat, we decided to subject plants to sequential treatments, with non-iron metals for 2 hours to trigger IRT1 ubiquitination followed by addition of biotin.

### BioID-based analyses of IRT1 proxitome

Since our goal was to better characterize IRT1’s function at the cell surface and the early events driving IRT1 internalization to TGN/EE, we sought to use iron-limited conditions combined to low non-iron metal concentrations for resupply (hereafter called -/sub) to limit IRT1 sorting to later endosomal compartments and vacuolar degradation. Plants were therefore grown and treated as described in Figure 2A. Briefly, plants were grown in absence of iron and non-iron metals, transferred to liquid growth conditions and exposed to low concentrations of non-iron metals for 2 hours before incubation with biotin. This experimental setup allowed to visualize a shared distribution of IRT1 between the plasma membrane and TGN/EE, with no impact on overall fluorescence level, and thus was chosen for Bio-ID (Supplemental Figure 2A-2C).

**Figure 2.**
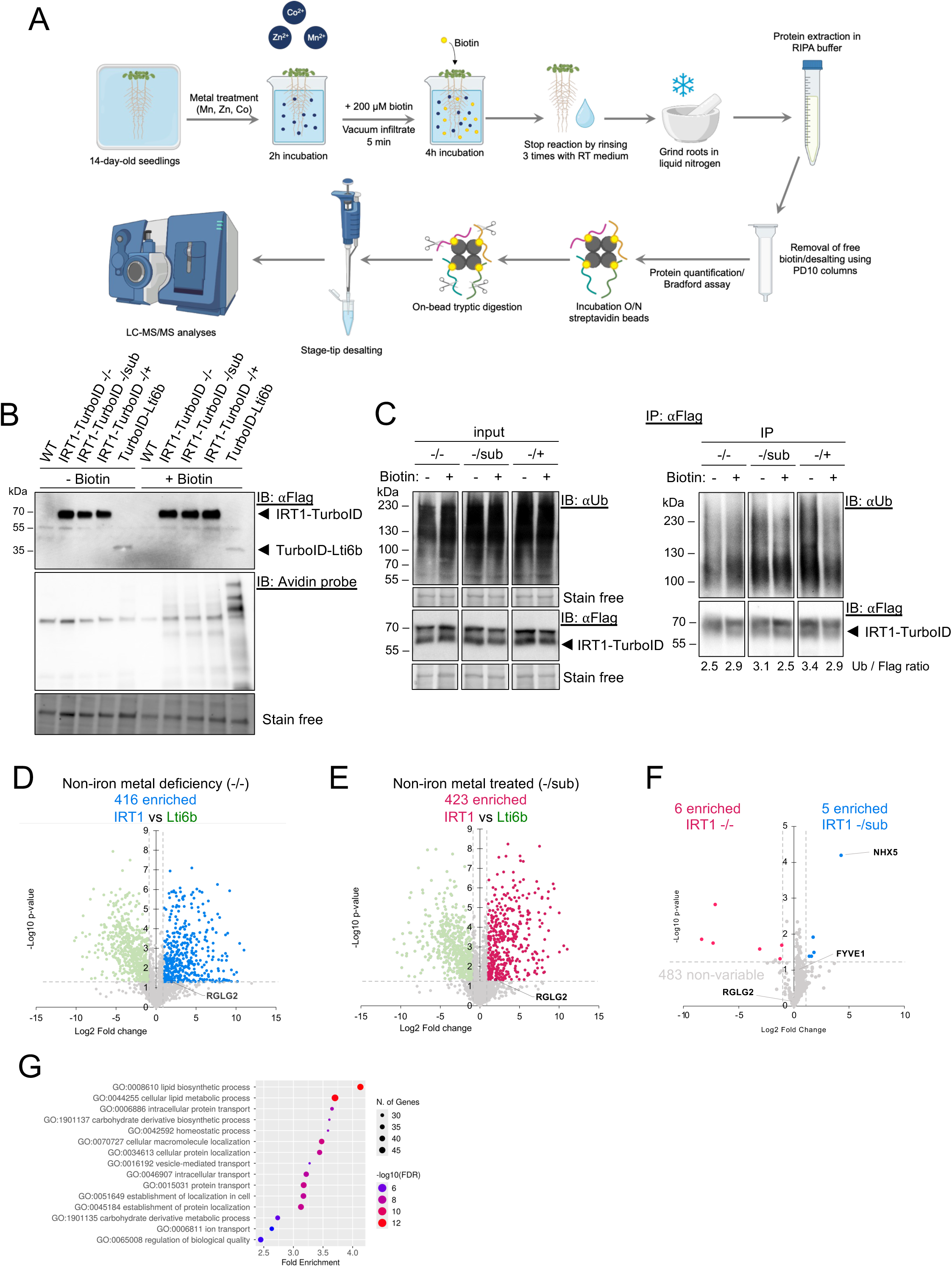
Identification of the IRT1 proximal proteome by TurboID. (A) Schematic representation of the experimental procedure, biotinylation treatment, streptavidin affinity purification and samples preparation for MS analyses. 14-day-old plants were treated with non-iron metal conditions (-/-, -/sub or -/+) for 2 h in liquid medium before 200 µM biotin was added for an additional 4 h. Streptavidin affinity purification was performed to recover biotinylated proteins before samples preparation for mass spectrometry analysis. (B) Protein levels of IRT1-TurboID and control TurboID-Lti6b fusions. 14-day-old plants grown as in (A) and roots were harvested for western blot analysis. TurboID fusion proteins and biotinylated proteins were detected using anti-Flag antibodies and the avidin probe, respectively. The stain free signal is used as loading control. (C-D) Volcan plots of differentially biotinylated proteins between IRT1-TurboID and the TurboID-Lti6b under control growth conditions (-/-) (C), and between IRT1-TurboID and the TurboID-Lti6b after a short term ressuply of non-iron metals (-/sub) (D). Proteins specifically enriched in IRT1 proxitome, compared to Lti6b, are shown in red (C) and blue (D), respectively. (E) Volcano plot of IRT1-specific proximal proteins extracted from (C) and (D). Proteins enriched in (-/-) or in (-/sub) are shown in red and blue, respectively. Proteins shown in grey are found irrespective of non-iron metal status. (F) Gene ontology (GO) analyses of biological processes of IRT1-specific proximal proteins (386 proteins from (E)) as a function of the fold enrichment. The number of genes and - log10(FDR) are also displayed. (G) Alt Text : Illustration and data on the TurboID analyses carried out with IRT1 and leading to the identification of IRT1 proximal proteins.

IRT1-TurboID and control plants were grown as described above, and roots were collected before being subjected to protein extraction. Protein extracts were used for western blot analyses to confirm that i) IRT1-TurboID and corresponding biotinylated proteins accumulated to similar extent across the metal-deficient and metal resupplied conditions, and that IRT1-TurboID was still be ubiquitinated (Figure 2B, 2C). To identify IRT1 proximal proteins, we performed affinity purification of biotinylated proteins in four replicates. Biotinylated proteins bound to streptavidin beads were eluted using boiling SDS supplemented with biotin and analyzed by liquid chromatography-tandem mass spectrometry (LC-MS/MS). Our analysis identified a total of 3042 proteins (Supplemental Dataset 1). Among the identified proteins found in IRT1-TurboID proxitome and absent from the TurboID-Lti6b proxitome, 416 were considered IRT1-specific proximal proteins based on the threshold of p-value ≤ 0.05 and fold change (IRT1/Lti6b) ≥ 2 (Figure 2D ; Supplemental Dataset 2). Non-iron metal addition allowed the isolation of 423 IRT1-specific proximal proteins using the same criteria (Figure 2E ; Supplemental Dataset 3). Comparison between IRT1 proximal proteins in non-iron metal deficient and non-iron metal resupplied conditions identified 494 proteins enriched at least in one of the conditions compared to Lti6b (Figure 2F ; Supplemental Dataset 4), including 6 specifically enriched in non-iron metal-depleted conditions (Figure 2F ; Supplemental Dataset 5) and 5 enriched specifically in non-iron metal resupplied conditions (Figure 2F ; Supplemental Dataset 6). Among IRT1 proximal proteins was found the FYVE1 endosomal protein that was previously reported to interact with IRT1 and control intracellular sorting and polarity of IRT1 (Barberon et al., 2014), thus validating our BioID approach. Gene ontology (GO) analyses was carried out using all 494 IRT1-specific proximal proteins to highlight biological functions associated. Three main families of GO terms were enriched : lipid metabolism, protein localization and ion homeostasis (Figure 2G). Of particular interest to our understanding of the dynamics of IRT1 and its post-translational modification are several proteins related to the ubiquitination machinery including several putative E3 ligases, among which RGLG2, ubiquitin-binding proteins, or ubiquitin-specific proteases. RGLG2 is found in both non-iron metal deficient and non-iron metal resupplied proximal protein datasets, highlighting that non-iron metals do not impact on the proximity between IRT1 and RGLG2, at least to the ∼10nm resolution of TurboID. Interestingly, previous work suggested a connection between RGLG2 and iron homeostasis (Pan et al., 2015), making of RGLG2 an interesting candidate to follow up on. Several candidates from the IRT1-specific proximal protein dataset are also associated to ion homeostasis, including metal transporters from the ZIP and MTP families as well as enzymes involved in coumarin biosynthesis. Among transport proteins is also found the NHX5 Na^+^(K^+^)/H^+^ antiporter that stands out as the most enriched protein in the IRT1-specific proximal proteins upon non-iron metal resupply. NHX5, and its NHX6 counterpart, are localized to the TGN/EE and late endosomes and have been shown to control endosomal ion homeostasis and TGN/EE trafficking (Dragwidge et al., 2019).

### Validation of NHX5 and RGLG2 as IRT1 proximal proteins and interactors

To showcase the ability of BioID approaches of plasma membrane proteins to identify new partners and regulators, we decided to validate deeper the two proximal proteins candidates RGLG2 and NXH5. Validation included the following steps : evaluation of protein-protein interaction between IRT1 and RGLG2 or NHX5, overlapping expression territories in root epidermal cells, and colocalization at the plasma membrane or in endosomal compartments.

To evaluate the ability of IRT1 to interact with its RGLG2 and NHX5 proximal proteins, we took advantage of the recently reported ALFA tag/ALFA nanobody (NB) technology and ratiometric tripartite functional fluorescence complementation assay (TriFC) (Neveu et al., 2025). TriFC uses coexpression of the functional IRT1-ALFA fusion, the ALFA NB-mCitN fusion, and mCitC-tagged NHX5 or RGLG2 in *Nicotiana benthamiana*, allowing reconstitution of functional mCit upon interaction between IRT1 and NHX5 or RGLG2 (Supplemental Figure 3A). The IRT1:AHA2 interaction we previously uncovered (Martín-Barranco et al., 2020; Neveu et al., 2025) was used as positive control, and expression of the ALFA NB without mCitN as negative control. TriFC-based reconstitution of mCit was observed at the plasma membrane for IRT1 and NHX5 and in intracellular vesicles (Figure 3A, 3B), which are likely TGN/EE where both proteins have previously been found (Barberon et al., 2011; Dubeaux et al., 2018; Dragwidge et al., 2019). Ratiometric TriFC also allowed quantification of the reconstituted mCit fluorescence relative to constitutively expressed mCherry, and confirmed the IRT1:NHX5 interaction (Figure 3B). This suggests that both proteins are not only proximal but also interact in plant cells. The closely-related NHX6 Na^+^(K^+^)/H^+^ antiporter also interacted with IRT1 in TriFC approaches (Supplemental Figure 3B, 3C). Coexpression of TriFC components for IRT1 and RGLG2 also revealed mCit reconstitution at the plasma membrane, while no mCit signal could be observed for the ALFA NB negative control (Figure 3A, 3B).

**Figure 3.**
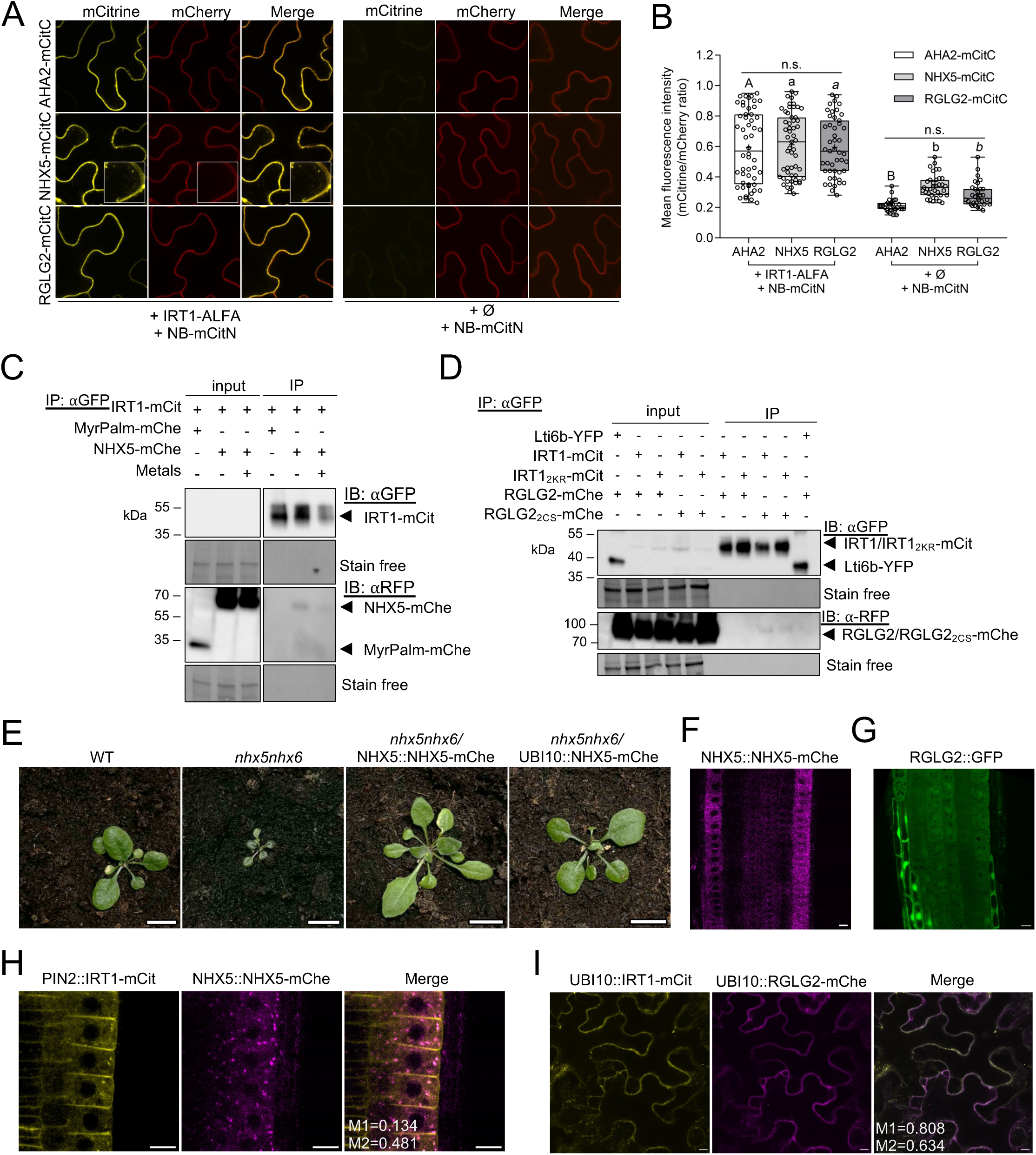
Validation of the candidate IRT1 proximal proteins NHX5 and RGLG2. (A) Trimolecular fluorescence complementation (TriFC) assay in *Nicotiana benthamiana* leaves coexpressing ALFA NB-mCitN with AHA2-mCitC (top, positive control), NHX5-mCitC (middle) or RGLG2-mCitC (bottom) in presence (left) or absence (Ø, right) of IRT1-ALFA. The absence of IRT1-ALFA (Ø) serves as negative control. The fluorescence emanating from the constitutively-expressed and cotransformed MyrPalm-mCherry plasma membrane marker serves as internal control for ratiometric quantification in (B). Scale bars, 10 μm. (B) Ratiometric quantification of the mCitrine/MyrPalm-mCherry fluorescence signal ratios from confocal microscopy images of *N. benthamiana* shown in (A). Box-and-whisker plots indicate the median (line), interquartile range (box), 1.5 interquartile range (whiskers), and mean values (‘+’ symbol). Experiments were done in triplicates where six cells from two independent leaves were imaged. Data were analyzed by two-way ANOVA followed by Bonferroni multiple comparison post-test. Statistically significant differences between combinations (presence *vs* absence of IRT1-ALFA) are indicated by letters (P<0.05). Tested interactions were statistically different from negative controls. No statistical difference (n.s.) was observed among all tested interactors. (C) Interaction of IRT1 with NHX5 by coimmunoprecipitation using transient expression in *N. benthamiana*. Protein extracts from leaves expressing IRT1-mCit together with NHX5-mChe were subjected to immunoprecipitation with anti-GFP antibodies. The plasma membrane-localized MyrPalm-mChe reporter was used as negative control. Coimmunoprecipitation of NHX5-mChe was detected using anti-RFP antibodies. The stain free signal is used as loading control for inputs. (D) Interaction of IRT1 with RGLG2 by coimmunoprecipitation using transient expression in *N. benthamiana*. Protein extracts from leaves expressing IRT1-mCit or IRT1_2KR_-mCit together with RGLG2-mChe or RGLG2_2CS_-mChe were subjected to immunoprecipitation with anti-GFP antibodies. Lti6b-YFP was used as negative control. Coimmunoprecipitation of RGLG2-mChe or RGLG2_2CS_-mChe was detected using anti-RFP antibodies. The stain free signal is used as loading control for inputs. (E) Phenotype of wild-type (WT), *nhx5nhx6*, *nhx5nhx6*/NHX5::NHX5-mChe, and *nhx5nhx6*/UBI10::NHX5-mChe. (F) Tissue localization of *NHX5* promoter activity. Confocal microscopy image of complemented plants expressing *nhx5nhx6*/NHX5::NHX5-mChe. (G) Tissue localization of *RGLG2* promoter. Confocal microscopy image of plants expressing GFP under the control of the *RGLG2* promoter. (H) Subcellular localization and colocalization of IRT1-mCit and NHX5-mChe. Confocal microscopy images of roots from stable transgenic plants coexpressing PIN2::IRT1mCit and NHX5::NHX5-mChe. The M1 (IRT1 overlapping with NHX5) and M2 Manders’ (NHX5 overlap with IRT1) colocalization coefficients are shown (I) Subcellular localization and colocalization of transiently-expressed RGLG2-mChe in *N. benthamiana* with IRT1-mCit. The M1 (IRT1 overlapping with RGLG2) and M2 Manders’ (RGLG2 overlap with IRT1) colocalization coefficients are shown. (J) Alt Text : Microscopy images and blots validating the interaction between two candidate proximal proteins and IRT1.

To back up TriFC protein:protein interaction evidence using an orthogonal assay, we performed coimmunoprecipitation using transient expression of IRT1-mCit and NHX5-mCherry (mChe) in *N. benthamiana*. NHX5-mChe coimmunoprecipitated with IRT1-mCit, while no signal could be observed for the plasma membrane MyrPalm-mChe negative control (Figure 3C). This confirms the ability of IRT1 and NHX5 to interact in plant cells. When challenged with non-iron metals, IRT1-mCit accumulated to lower levels but coimmunoprecipitated similar NHX5-mChe levels (Figure 3C), consistent with the enrichment observed upon non-iron metal treatment in our TurboID dataset. Similar experiments were carried out with IRT1-mCit and RGLG2-mChe to confirm their physical interaction in live cells using transient expression in *N. benthamiana*. No interaction between RGLG2 and IRT1 could be observed in this assay (Figure 3D). However, considering the notoriously weak and transient interaction between E3 ubiquitin ligases and their targets, we decided to test the ability of mutant versions of IRT1 and RGLG2 aiming at stabilizing such interactions. To this purpose, we used the non-ubiquitinatable IRT1_2KR_ (Barberon et al., 2011; Dubeaux et al., 2018) and the RGLG2_2CS_ zinc finger mutant impaired in Zn^2+^ coordination to abolish ubiquitination (Garcia-Barcena et al., 2020). IRT1_2KR_-mCit failed to interact with RGLG2-mChe (Figure 3D). However, RGLG2_2CS_-mChe interacted with both IRT1-mCit and IRT1_2KR_-mCit, supporting further the ability of IRT1 and RGLG2 to partner in live cells and stressing the edge of BioID approaches to pick up weak transient interactions.

Since both TriFC and coimmunoprecipitation protein:protein interaction evidences are based on overexpression, we next ascertained that both *NHX5* and *RGLG2* are found at least in root epidermal cells to support their capacity to physically interact with IRT1. We generated a complemented *nhx5nhx6*/NHX5::NHX5-mCherry line that showed a wild-type phenotype, confirming the functionality of the NHX5-mChe fusion (Figure 3E). NHX5-mChe driven by the *NHX5* endogenous promoter was observed in the root epidermis (Figure 3F), confirming that NHX5 and IRT1 coexist in similar cell types. In parallel, *RGLG2* expression territories were assessed by generating a transcriptional reporter line expressing mCit under the control of the *RGLG2* promoter. *RGLG2* promoter activity was observed across all root cell types (Figure 3G), suggesting that *RGLG2* shares common expression territories with *IRT1* and further strengthening the protein interaction data obtained from TurboID, TriFC and coimmunoprecipitation.

NHX5 is localized to the TGN/EE (Dragwidge et al., 2019), where IRT1 transits through during endocytosis (Barberon et al., 2011; Dubeaux et al., 2018). We confirmed their partial subcellular overlap using transgenic expressing both PIN2::IRT1-mCit and UBI10::NHX5-mChe, where intracellular vesicles positive for both IRT1 and NHX5 are observed (Figure 3H). RGLG2 was previously reported to be myristoylated and associated to the plasma membrane (Yin et al., 2007). We evaluated the subcellular localization of RGLG2 using transient expression of a RGLG2-mChe fusion in *N. benthamiana* and confirmed that, consistent with the literature, RGLG2 is mostly found at the plasma membrane where it colocalizes with IRT1-mCit (Figure 3I).

Altogether, the evidence gathered here confirmed the ability of IRT1 to interact with RGLG2 and NHX5 and their presence in overlapping territories and subcellular localizations.

### NHX5 and NHX6 contribute to plant non-iron metal responses

To better characterize the relationship between NHX5, IRT1 and responses to non-iron metals, we first assessed the colocalization extent of PIN2::IRT1-mCit and UBI10::NHX5-mChe in stable transgenic lines exposed to various non-iron metal provision. IRT1-mCit showed increased internalization from the plasma membrane to TGN/EE with rising concentrations of non-iron metals (-/- to -/+), as expected (Dubeaux et al., 2018). NHX5-mChe displayed a punctate pattern consistent with its presence in the TGN/EE (Dragwidge et al., 2019). In agreement with the heightened localization of IRT1-mCit in TGN/EE, its colocalization with the TGN/EE-localized NHX5-mChe increased with non-iron metals, with a maximum observed in -/+ conditions (Figure 4A, 4B).

**Figure 4.**
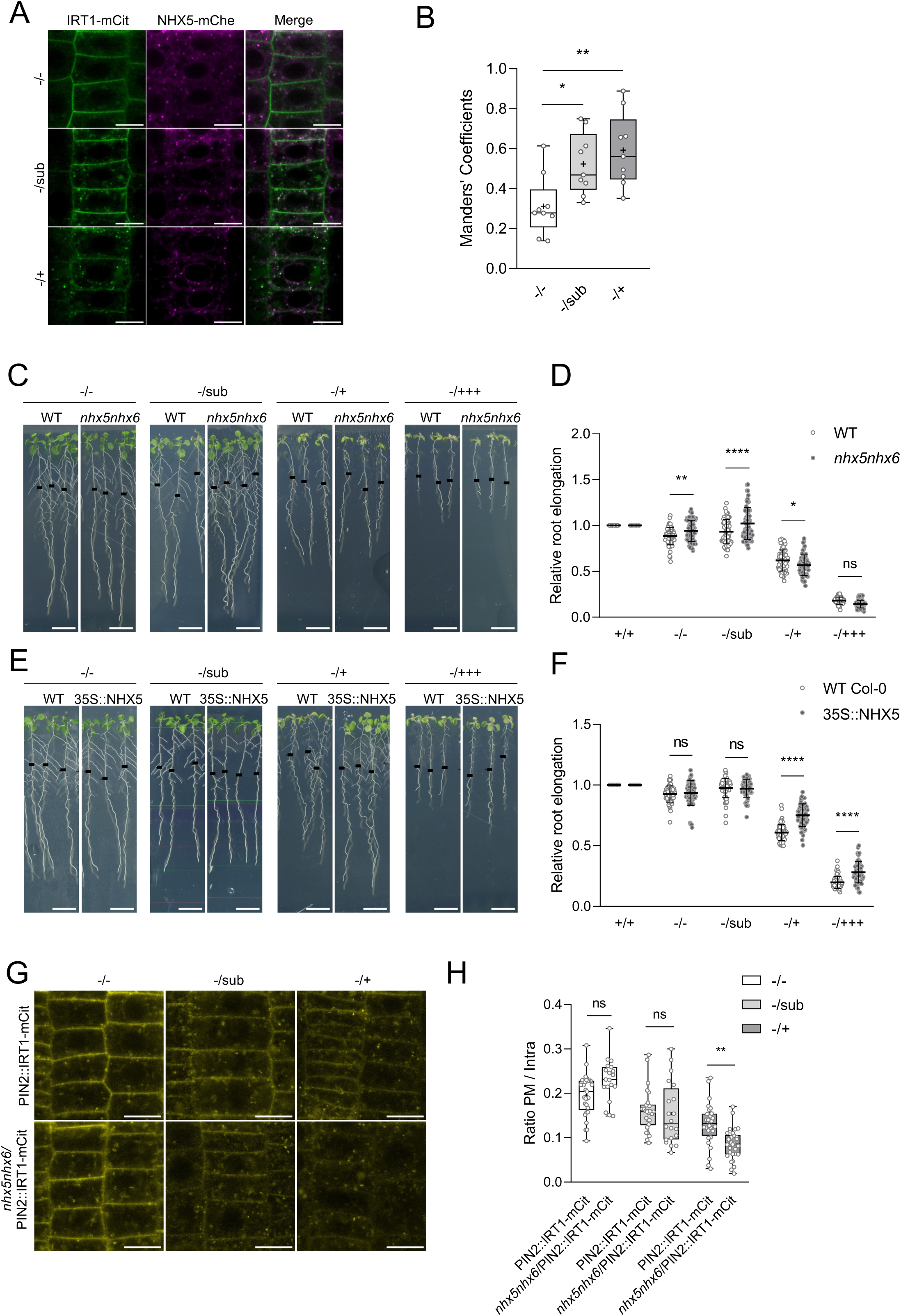
NHX5 and NHX6 participate to plant metal nutrition. (A) Impact of non-iron metal provision on the colocalization of NHX5 and IRT1. Confocal microscopy images of plants coexpressing PIN2::IRT1-mCit and UBI10::NHX5-mChe treated various non-iron metal provision( -/-, -/sub and -/+) for 3 h. Scale bar, 10 μm. (B) Quantification of the colocalization of NHX5 with IRT1 shown in (A). Analysis was performed using ImageJ JACoP plugin. Manders’ colocalization coefficients (NHX5 overlap with IRT1) were plotted. Experiments were done in triplicates where three cells from three independent roots were imaged. Error bars represent standard deviation. Asterisks indicate significant (one-way ANOVA, Bonferroni’s multiple comparisons test, **P < 0.01; *P < 0.05; ns, not significant). (C) Root length phenotyping analysis of *nhx5nhx6* in different non-iron metal conditions. Wild-type (WT) and n*hx5nhx6* mutant plants were grown for 7 days on +/+ before being transferred for 7 days on the different metal conditions. Black lines shows the root length at the time of transfer. Scale bar, 1 cm. (D) Quantification of root elongation of wild-type (WT) and n*hx5nhx6* mutant plants grown as in (C). Data are presented as ratio to the +/+ condition where *IRT1* is not expressed. Experiments were done in triplicates (n=24). Error bars represent standard deviation. Asterisks indicate significant (two-way ANOVA , Sidakpost hoc test, ****P < 0.0001; **P < 0.01; *P < 0.05; ns, not significant). (E) Root length phenotyping analysis of *nhx5nhx6*/35S::NHX5-YFP grown as in (C). Black lines shows the root length at the time of transfer. Scale bar, 1 cm. (F) Quantification of root elongation wild-type (WT) and *nhx5nhx6*/35S::NHX5-YFP. Data are presented as ratio to the +/+ condition where *IRT1* is not expressed. Experiments were done in triplicates (n=24). Error bars represent standard deviation. Asterisks indicate significant (two-way ANOVA , Sidakpost hoc test, ****P < 0.0001 ; ns, not significant). (G) Role of NHX5 and NHX6 in the non-iron metal-induced endocytosis of IRT1. Confocal microscopy images of plants coexpressing PIN2::IRT1-mCit and *nhx5nhx*6/PIN2::IRT1-mCit treated various non-iron metal provision ( -/-, -/sub and -/+) for 3 h. Scale bar, 10 μm. (H) Quantification of the plasma membrane to intracellular fluorescence ratio of plants coexpressing PIN2::IRT1-mCit and *nhx5nhx*6/PIN2::IRT1-mCit grown as in (G). Experiments were done in duplicates where five cells from three independent roots were imaged. Error bars represent standard deviation. Asterisks indicate significant (one-way ANOVA, Bonferroni’s multiple comparisons test, **P < 0.01 ; ns, not significant). Alt Text : Microscopy data phenotypic analyses characterizing the role of NHX5 and NHX6 as new IRT1 regulators.

Plants grown under iron starvation, thus expressing high levels of *IRT1*, and exposed to non-iron metal excess have previously been shown to show reduced root growth (Dubeaux et al., 2018). To evaluate the genetic contribution of *NHX5*, and its *NHX6* redundant counterpart, in plant responses to non-iron metals, we characterized root growth responses of the *nhx5nhx6* double loss-of-function mutant (Bassil et al., 2011). To this purpose, wild-type and *nhx5nhx6* plants were grown for 7 days in standard half-strength Murashige and Skoog medium (MS/2, (Murashige and Skoog, 1962), containing iron and non-iron metals (+/+) before being transferred to plates lacking iron and non-iron metals (-/-), lacking iron and containing low concentrations of non-iron metals (-/sub, as used for the BioID), lacking iron and containing half MS levels of non-iron metals (-/+), or lacking iron and containing non-iron metals excess (-/+++ as previously reported in (Dubeaux et al., 2018) (Figure 4C). Root growth after transfer was measured and scored relative to control conditions (+/+), highlighting how *nhx5nhx6* responds to non-iron metals compared to wild-type plants. Under conditions where *IRT1* is expressed and the corresponding protein is mostly found at the plasma membrane (-/- and - /sub), *nhx5nhx6* appears to be slightly less impacted than wild-type (Figure 4C, 4D). Increasing non-iron metal concentrations, conditions where IRT1 is less at the plasma membrane and is targeted to endosomal compartments (-/+) and the vacuole (-/+++), yielded a slight non-iron metal hypersensitivity (Figure 4C, 4D). Therefore NHX5 and NHX6 function seems to be conditional and differ between non-iron metal provision and associated IRT1 localization.

Considering that the observed effects with *nhx5nhx*6 are mild and that the *NHX* gene family in Arabidopsis contains eight members, we conducted a gain-of-function approach using the previously reported *nhx5nhx6*/35S::NHX5-YFP line (Bassil et al., 2011) to better characterize the involvement of NHX-type antiporters in plant responses to non-iron metals. *NHX5* overexpression yielded increased resistance to non-iron metals in conditions where IRT1 is gradually removed from the cell surface (-/+) and degraded in the vacuole (-/+++) (Figure 4E, 4F), highlighting further the negative genetic relationship between NHX-type antiporters and responses to non-iron metals.

NHX5 and NHX6 were previously shown to be required for endosomal trafficking and recycling of endocytosed BRI1 to the cell surface, while not necessary for the recycling of PIN1 and PIN2 (Dragwidge et al., 2018; Dragwidge et al., 2019). To further examine the role of NHX-type antiporters in responses to non-iron metals, we monitored the well-established endocytosis and degradation of IRT1 triggered by non-iron metals (Dubeaux et al., 2018). We therefore monitored IRT1-mCit localization driven by the *PIN2* promoter in wild-type and in *nhx5nhx6 mutant* in response to various non-iron metal provision*. PIN2* promoter was chosen to drive IRT1-mCit expression to i) obtain constitutive IRT1-mCit expression and thus focus on post-transcriptional events, and ii) image IRT1 endocytosis in epidermal cells at the root tip that are well suited to track the precise localization of plasma membrane proteins. Our previous work demonstrated that IRT1 undergoes metal-dependent endocytosis regardless of the promoter used (Dubeaux et al., 2018; Spielmann et al., 2022). Surprisingly, we observed that the metal-induced degradation of IRT1 was enhanced in *nhx5nhx6*, highlighted by the lower plasma membrane:intracellular fluorescence ratio (Figure 5G, 5H). Although this observation may be counterintuitive considering previous reports on the influence of NHX5 and NH6 on endocytosis, it may reflect a role of endosomal pH on endosomal protein recycling (Dragwidge et al., 2018) or on endosomal metal transport from the TGN/EE to the cytoplasm.

**Figure 5.**
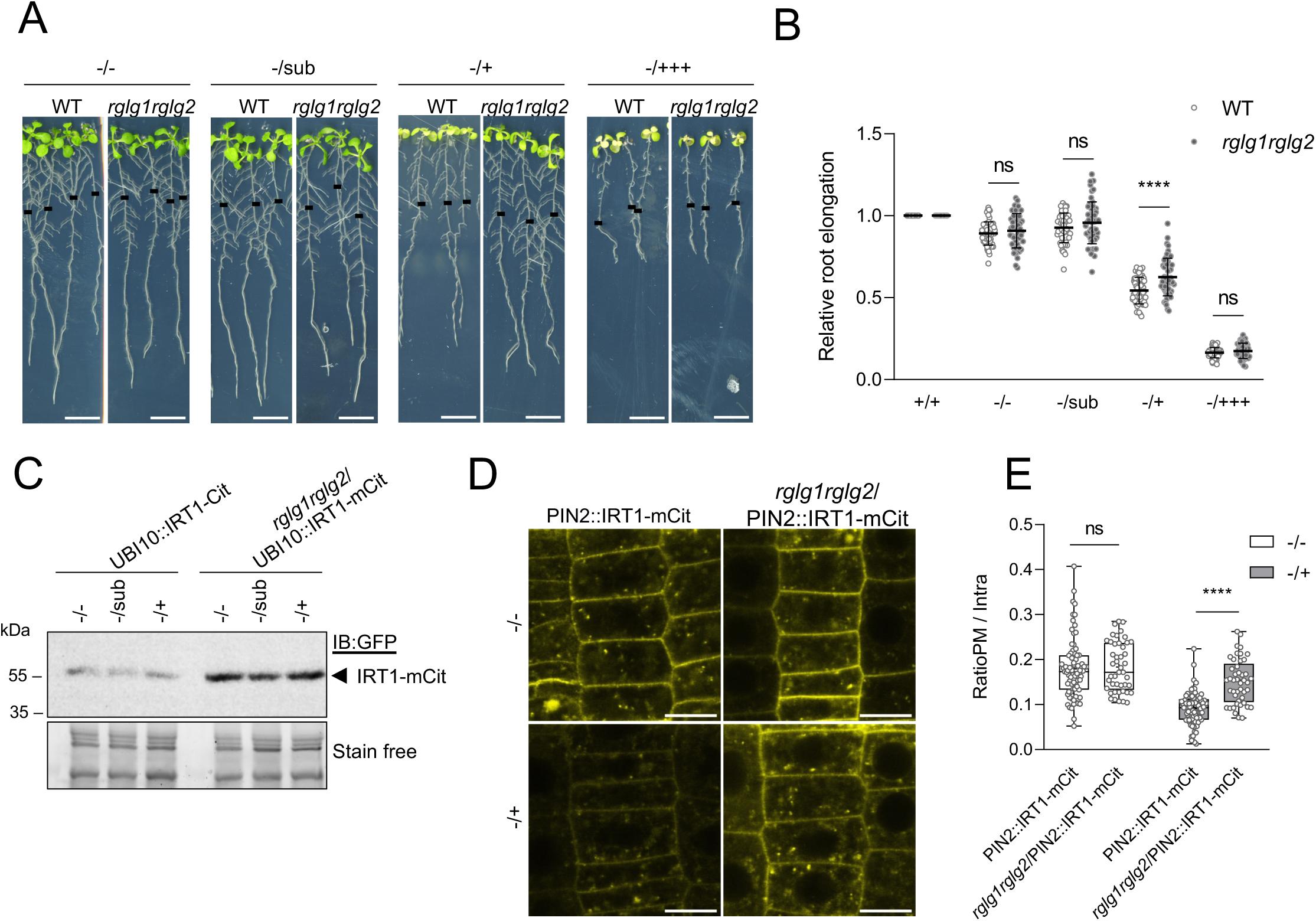
RGLG2 regulates IRT1 levels and plasma membrane localization. (A) Root length phenotyping analysis of *rglg1rglg2* in different non-iron metal conditions. Wild-type (WT) and *rglg1rglg2* mutant plants were grown for 7 days on +/+ before being transferred for 7 days on the different metal conditions. Black lines shows the root length at the time of transfer. Scale bar, 1 cm. (B) Quantification of root elongation of wild-type (WT) and *rglg1rglg2* mutant plants grown as in (A). Data are presented as ratio to the +/+ condition where *IRT1* is not expressed. Experiments were done in triplicates (n=24). Error bars represent standard deviation. Asterisks indicate significant (two-way ANOVA , Sidakpost hoc test, ****P < 0.0001; **P < 0.01 ; ns, not significant). (C) Impact of the loss of RGLG1 and RGLG2 on IRT1 protein accumulation. Plants expressing UBI10::IRT1-mCit or *rglg1rglg2*/UBI10-IRT1-mCit were treated with various non-iron metal provision and IRT1-mCit protein accumulation in root monitored by western blot using anti-GFP antibodies. The stain free signal is used as loading control. (D) Role of RGLG1 and RGLG2 on the non-iron metal-induced endocytosis or IRT1. Confocal microscopy images of plants coexpressing PIN2::IRT1-mCit and *rglg1rglg2*/PIN2::IRT1-mCit treated various non-iron metal provision ( -/-, -/sub and -/+) for 3 h. Scale bar, 10 μm. (E) Quantification of the plasma membrane to intracellular fluorescence ratio of plants coexpressing PIN2::IRT1-mCit and *rglg1rglg2*/PIN2::IRT1-mCit grown as in (D). Experiments were done in triplicates where five cells from three independent roots were imaged. Error bars represent standard deviation. Asterisks indicate significant (one-way ANOVA, Bonferroni’s multiple comparisons test, ****P <0.0001 ; ns, not significant). Alt Text : Microscopy data and phenotypic analyses characterizing the role of RGLG2 and RGLG1 as new IRT1 regulators.

### RGLG2 regulates non-iron metal responses and IRT1 endocytosis

Considering the reported redundancy of RGLG2 with RGLG1, we used the previously reported the *rglg1rglg2* double mutant (Yin et al., 2007) to evaluate its response to non-iron metal provision. Root elongation of wild-type and *rglg1rglg2* after transfer from +/+ to iron deficient conditions, combined with various non-iron metal regimes, was scored. *rglg1rglg2* appeared overall less impacted by non-iron metal provision than wild-type plants (Figure 5A, 5B). This suggests that genetically, RGLG1 and RGLG2 act as negative regulators of plant non-iron metal responses. Since RGLG2 was found in the proximal proteome of IRT1 and to interact with IRT1, we first sought to evaluate the impact of RGLG2 transient overexpression in *N. benthamiana* on IRT1 endocytosis in response to non-iron metals. As controls, we used the non-ubiquitinatable IRT1_2KR_ and the ubiquitination-defective RGLG2_2CS_. In the absence of non-iron metal treatment (i.e. standard conditions), coexpression of IRT1_2KR_ and RGLG2, IRT1 and RGLG2_2CS_, and IRT1_2KR_ with RGLG2_2CS_ all yielded a plasma membrane localization of IRT1 and no or little IRT1 endocytosis using endosome counts as proxy (Supplemental Figure 4A, 4B). In sharp contrast, coexpression of IRT1 and RGLG2 greatly reduced plasma membrane levels of IRT1 and increased endosome numbers (Supplemental Figure 4A, B), indicating that RGLG2 overexpression promotes IRT1 endocytosis. Treatment of *N. benthamiana* with non-iron metals was previously shown to promote IRT1 endocytosis (Cointry et al., 2025). Consistently, IRT1 endocytosis was enhanced upon non-iron metal exposure when coexpressed with the ubiquitination-defective RGLG2_2CS_ (Supplemental Figure 4A, 4B). This illustrates the endogenous non-iron mechanism present in *N. benthamiana* leaves. Coexpression of IRT1 and RGLG2 did not show greater endocytosis of IRT1 upon non-iron metal treatment compared to standard conditions, likely because RGLG2 overexpression already depleted IRT1 from the plasma membrane. Consistent with previous reports in Arabidopsis (Dubeaux et al., 2018), the ubiquitination-defective IRT1_2KR_ variant failed to respond to metals. Altogether, this clearly highlights that overexpression of RGLG2 in N. benthamiana impinges on IRT1 endocytosis and levels. To confirm these observations in Arabidopsis, IRT1 protein levels were monitored in wild-type and *rglg1rglg2*. To this purpose, we generated the *rglg1rglg2*/UBI10::IRT1-mCit line constitutively expressing IRT1-Cit (Spielmann et al., 2022) in the *rglg1rglg2* mutant background. In all non-iron metal conditions tested, *rglg1rglg2* accumulated higher levels of IRT1-mCit protein compared to wild-type plants (Figure 5C). This confirms the existence of a post-transcriptional regulation of *IRT1* by RGLG2 in *Arabidopsis* plants. We validated further these observations by imaging PIN2::IRT1-mCit in wild-type and *rglg1rglg2*. Metal provision lowered IRT1-mCit plasma membrane levels in the wild-type background, as visualized by the decreased ratio between plasma membrane and intracellular fluorescence (Figure 5D, 5E). In contrast, IRT1-mCit significantly remained more at the cell surface upon non-iron metal exposure in *rglg1rglg2,* indicating that the RGLG1 and RGLG2 E3 ubiquitin ligases directly regulate the metal-dependent and ubiquitin-mediated endocytosis of IRT1.

## Discussion

The main root iron transport machinery components and their associated regulatory networks were initially identified in Arabidopsis and non-grasses plants by forward genetic screens looking for mutant with altered macroscopic or molecular phenotypes (Yi and Guerinot, 1996; Ling et al., 2002; Colangelo and Guerinot, 2004), through reverse genetic approaches screening for phenotypes of interest (Shin et al., 2013), or using homology with non-plant organisms (Eide et al., 1996; Curie et al., 2000; Thomine et al., 2000; Ivanov et al., 2014). The search for interacting proteins based on yeast two hybrid screening, or directed yeast two hybrid tests using candidate partners, and AP-MS allowed a rapid expansion of our knowledge about the transcriptional networks and post-translational regulatory mechanisms controlling iron transporters, and especially for IRT1 and NRAMP1 (Barberon et al., 2011; Barberon et al., 2014; Dubeaux et al., 2018; Martín-Barranco et al., 2020; Fu et al., 2022). Here, we unveil new regulators of plant metal uptake by pioneering proximity labelling strategy for the IRT1 metal transporter.

Proximity labelling and TurboID approaches have been developed to identify proteins proximal to a target of interest within a certain radius in living cells, providing new insights into protein-protein interactions and subcellular localization. Most importantly, the stable labeling of proximal proteins holds the premise of picking up transient or weak interacting partners compared to traditional AP-MS strategies, allowing the exploration of a new repertoire of regulators that was previously hardly accessible. Not surprisingly, the IRT1 proximal protein showed only minor overlap with the previously reported characterization of the IRT1 complex composition by AP-MS (Martín-Barranco et al., 2020) (Supplemental Figure S5), keeping in mind that plant growth conditions used slightly differ between the two datasets. Regardless, that clearly illustrates the need for several orthogonal assays to draw the most comprehensive picture of the interaction networks for a protein of interest. It is noteworthy that the proteins specifically found in the IRT1 proxitome and absent from the AP-MS-based IRT1 complex contain many signaling proteins such as kinases or E3 ubiquitin ligase (Supplemental Datasets 7-9), which are known to interact only weakly or transiently with their targets. Such observations also supports the edge of proximity labeling strategies compared to AP-MS. Among IRT1 proximal proteins, the NHX5 Na^+^(K^+^)/H^+^ antiporter and the RGLG2 E3 ubiquitin ligase were selected for further analyses because they either represent proteins unrelated to metal nutrition (NHX5), or proteins that are difficult to identify using other protein:protein interaction strategies (RGLG2). For both, their ability to interact with IRT1 was confirmed using TriFC, coimmunoprecipitation, and their presence in overlapping expression territories and subcellular compartments with IRT1 was validated. Reverse genetic approaches using knock-out mutants revealed their respective involvement in plant metal nutrition and/or responses to non-iron metals. The fact that macroscopic phenotypes observed for *nhx5nhx6* and *rglg1rglg2* are rather subtle suggests that additional members of the corresponding families are likely playing a role in IRT1-mediated metal nutrition as well. This is supported by the stronger gain-of-function phenotype obtained with *NHX5* overexpression, for example. These observations also argue for the ability of TurboID to identify proteins that play minor roles in biological processes, and that can hardly be identified through classical genetic approaches due to the absence of strong phenotypes.

The contribution of NHX5 and NHX6 to plant responses to non-iron metals appears to be complex and depends on non-iron metal provision, with a positive role observed for low concentrations and a negative role at elevated concentrations. The use of *NHX5* overexpressor confirms the implication of NHX-type antiporters to non-iron metal nutrition and points to their negative role. NHX5 and NHX6 were previously shown to impact on endosomal trafficking in a cargo-specific manner (Dragwidge et al., 2018; Dragwidge et al., 2019). Loss of *NHX5* and *NHX6* revealed a faster endocytosis and degradation of IRT1 upon non-iron treatment. This is reminiscent of the reduced PIN1 and PIN2 protein levels observed in *nhx5nhx6* (Dragwidge et al., 2018). The fact that the cytosolic tail of NHX6 interacts with the retromer complex protein SNX1 (Ashnest et al., 2015), associated to the lower PIN abundance in *nhx5nhx6*, led to the assumption that NHX5 and NHX6 may assist retromer-mediated retrieval of PINs. This would be consistent with the increased endocytosis of IRT1 observed in response to non-iron metals. However, such scenario is not consistent with the observed phenotypes derived from *nhx5nhx6* or *NHX5* overexpressor plants. Considering that NHX5 and NHX6 likely impact on the trafficking of many plasma membrane proteins, determining the functional outcome of *NHX5* and *NHX6* loss is difficult. In addition, NHX5 and NHX6 mediate endosomal ion and pH homeostasis, with endosomal compartments being more acidic in *nhx5nhx6* (Reguera et al., 2015). At the plasma membrane, IRT1-mediated metal uptake requires H^+^ export by the IRT1 complex member AHA2 (Santi and Schmidt, 2009; Martín-Barranco et al., 2020). One can speculate that IRT1 may also drive metal import into the cytosol from TGN/EE and that acidification by the TGN/EE-localized vATPase and H^+^ leak driven by NHX5 and NHX6 likely impact on plant metal nutrition. Future work evaluating pH in TGN/EE pH in response to non-iron metals and the contribution of TGN/EE vATPase and NHX5/NHX6 will be necessary to better understand the phenotype of *nhx5nhx6*.

Wild-type plants harbor bifurcated root hairs in response to iron starvation. r*glg1rglg2* was found to show constitutively high number of branched root hairs (Li and Schmidt, 2010). This points to a role of RGLG1 and RGLG2 in developmental responses to iron, although the underlying molecular mechanisms are unknown. The identification of RGLG2 as an IRT1 proximal protein and partner regulating IRT1 protein levels and endocytosis raises new questions as to the biological function of RGLG2 and its close homolog RGLG1. The reduced phenotypic sensitivity of *rglg1rglg2* to non-iron metal provision contrasts with the reduced endocytosis of IRT1. RGLGs likely target diverse proteins in plant cells, and act both in developmental (branched root hairs) and cellular (IRT1 endocytosis) responses to metals. This greatly complicates interpretation of macroscopic phenotypes that result from many different input responses. Undoubtedly, RGLGs regulate IRT1 protein abundance and cell surface levels. To date, IDF1 was the only E3 ubiquitin ligase known to regulate IRT1 (Shin et al., 2013). IDF1 drives the ubiquitin-mediated endocytosis and vacuolar degradation of IRT1 and thus contributes to iron and non-iron metal homeostasis (Dubeaux et al., 2018). RGLG1 and RGLG2 are unlikely to act in a separate pathway from the ubiquitin-mediated degradation of IRT1 in response to non-iron metals since *rglg1rglg2* is hypersensitive for the non-iron metal-induced endocytosis of IRT1 readout. Rather, RGLGs may act redundantly with the unrelated IDF1 E3 ligase to fine tune IRT1 levels. The rationale for having different E3s controlling IRT1 in response to the same cue is unclear, but may reveal time- or developmental stage-specific ubiquitination of IRT1 in response to non-iron metals. Alternatively, RGLGs and IDF1 may act sequentially during IRT1 ubiquitination. IRT1 was demonstrated to undergo K63 polyubiquitination in response to non-iron metals (Dubeaux et al., 2018). The formation of K63-linked polyubiquitin chains is under the control of the two UBC13-type E2 ubiquitin-conjugating enzymes, UBC35 and UBC36 in Arabidopsis (Romero-Barrios et al., 2020). Structural analyses of the human K63 polyubiquitination machinery containing UBC13, the UEV1 E2 variant, and ubiquitin revealed interesting insight into K63-linked chain formation. The active site cysteine of UBC13 is covalently bound to a first donor Ub, while UEV1A binds non-covalently to a second acceptor Ub, presenting residue K63 for chain assembly (Hodge et al., 2016). This suggests that a priming monoubiquitination on a substrate protein is necessary for UBC13 to catalyze K63-linked chain extension (Soss et al., 2011; Mattiroli et al., 2012). RGLGs and IDF1 may therefore act at different steps along IRT1 ubiquitination, with RGLG2 possibly catalyzing the priming IRT1 ubiquitination while IDF1 would polymerize K63 polyubiquitin chains to drive endosomal sorting and vacuolar degradation. The fact that IDF1 was not identified among IRT1 proximal proteins under conditions where internalization is promoted but not vacuolar targeting is consistent with this view. Regardless, this hypothesis will necessitate deep in vitro biochemical investigation of RGLGs, IDF1 and their cognate E2s to decipher their exact role.

In summary, our study identified numerous novel interactors of the IRT1 transporter, underscoring the central role of IRT1 in upholding metal homeostasis to sustain optimal plant growth and development. Unraveling the function of such IRT1 partners will enhance our understanding of integrated plant responses to metals. Our work on IRT1 proximity labeling also paves the way to applying TurboID to other transporters and channels, for which biochemical properties are still considered as a hurdle. This will be necessary to further our global comprehension of transporter regulation and plant nutrition.

## Methods

### Plant material and constructs

All experiments were performed on *Arabidopsis thaliana* ecotype Col-0. The *irt1* knockout mutant for the *IRT1* gene (AT4G19690) in the Col-0 background was generated by the CRISPR/Cas9 technique (Cointry et al., 2025). Knockout T-DNA insertion mutant lines were obtained from the Eurasian Arabidopsis Stock Centre (uNASC). The IRT1::IRT1-mCit, PIN2::IRT1-mCit and UBI10::IRT1-mCit expressing lines used in this study were previously characterized (Dubeaux et al., 2018; Spielmann et al., 2022). T-DNA knock-out mutant alleles used in this work were WiscDsLox345-348M8 (*nhx5-1*), double *nhx5nhx6* mutant (carrying *nhx5-2* (GABI_094H05) and *nhx6-2* (SALK_100042) alleles respectively), as well as the complemented line *nhx5nhx6*/35S::NHX5-YFP kindly donated by Elias Bassil (Bassil et al., 2011). The *rglg1rglg2* double mutant, based on *rglg1* (SALK_011892) and *rglg2* (SALK_062384) mutants, was kindly donated by Andreas Bachmair.

The TurboID-FLAG fusion was inserted into IRT1 coding sequence between residues S73 and R74 in pDONR221 using primers listed in Supplemental Table 1. The open reading frames of *NHX5* (AT1G54370), *RGLG2* (AT5G14420), and TurboID-FLAG fusion were cloned into pDONR221 Gateway entry vector, the open reading frame of Lti6b (AT3G05890) was cloned into pDONR2R-P3. Respective promotor regions and 5’UTRs (∼2500 nucleotides upstream of the coding sequence) were selected and cloned into pDONRP4-P1R entry vector using primers listed in Supplemental Table 1. Other pDONRP4-P1R vectors carrying *IRT1*, *UBI10* and *35S* promoters were previously described (Marquès-Bueno et al., 2016). pDONRP2R-P3 vectors carrying GFP, mCit and mChe were previously described (Marquès-Bueno et al., 2016). Final destination vectors were obtained by using MultiSite Gateway Three Fragment recombination system (Thermo), using pDONR vectors and gateway destination vectors (Karimi et al., 2007) to generate the following expression constructs : IRT1::IRT1-TurboID, UBI10::NHX5-mCherry, NHX5::NHX5-mCherry, RGLG2::GFP, UBI10::RGLG2-mCherry, and UBI10::RGLG2_CS_-mCherry. The list of constructs generated or used is listed in Supplemental Table 2.

Generated constructs were transformed into wild-type Arabidopsis by floral dipping or used in transient expression assays in *N. benthamiana*. For all stable plant constructs, independent lines homozygous for the transgene were selected in T3. Confocal microscopy, phenotypic analysis and protein extraction were performed on representative mono-insertional homozygous T3 lines. Introgression of tagged constructs into genotypes of interest was done by crossing and selection of homozygous progeny.

To generate the final constructs for TriFC experiments, pDONR221 plasmids carrying the proteins of interest or ALFA NB (Neveu et al., 2025) were recombined with pDONR-P2RP3 plasmids containing the N-terminal (mCitN) or C-terminal (mCitC) fragments of mCit (residues 1-155 and 155-238, respectively ; (Dolde et al., 2023) and with the pDONR P4P1R-UBI10 promoter or pDONR P4P1R-35S plasmids (Marquès-Bueno et al., 2016), into the pK7m34GW or pH7m34GW pDEST vectors (Karimi et al., 2007).

### Plant growth conditions

Following sterilization, seeds were placed at 4°C in the dark for 2 to 3 days for stratification before growing vertically under long day photoperiod (16 hours light/8 hours dark at 21°C). For biochemical experiments, plants were grown on solid half-strength MS medium lacking Fe and non-iron metals (-/-). After 14 days, plants were transferred for 5 hours to Fe-deficient liquid MS/2 medium supplemented with different metal concentrations (-/-, -/sub and -/+). -/+ corresponds to medium lacking Fe and containing standard non-iron metal concentrations as defined in half-strength MS medium, and -/sub corresponds to medium lacking Fe and containing suboptimal non-iron metal concentrations defined as ten times lower than the ones found in half-strength MS medium.

For BioID experiments, plants were seeded on MS/2 solid medium lacking Fe and non-iron metals (-/-). After 14 days, plants were transferred for 2 hours in liquid -Fe MS/2 supplemented with different metal concentrations (-/- ; -/sub) followed by biotin additional for another 4 hours (Figure 2A).

For confocal microscopy imaging, plants were grown on Fe- and metal-deficient (-/-) MS/2 solid medium for 7 days, then transferred for 3 hours to liquid MS/2 -Fe medium supplemented with different metal concentrations (-/- ; -/sub ; -/+).

For phenotyping, plants were grown on Fe-sufficient and standard metal supply (+/+) MS/2 solid medium for 7 days before being transferred to MS/2 solid -Fe-deficient medium supplemented with different metal concentrations (-/- ; -/sub ; -/+) for an additional 7 days.

### Affinity purification of biotinylated proteins

Biotinylation and subsequent streptavidin-based affinity purification protocol were done as previously described (Mair et al., 2019). *irt1*/IRT1::IRT1-TurboID and the IRT1::TurboID-Lti6b negative control were grown and treated as described above. After incubation with biotin, plants were washed three times with liquid MS/2 medium lacking iron and biotin to stop the reaction, and roots were harvested (Figure 2A). Roots were ground to fine powder in liquid nitrogen. 1 g of roots was resuspended in 2 mL of RIPA lysis buffer (50 mM Tris-HCl pH 7.5, 500 mM NaCl, 1% NP-40, 0.4% SDS, 0.5% sodium deoxycholate, 1 mM DTT, 1/100 protease inhibitor cocktail). Samples were then centrifuged twice at 3,800 g at 4°C for 10 minutes, before being placed on rotor wheel at 4°C for 1 hour and subjected to another centrifugation for 10 minutes at 10 000 g. PD-10 desalting columns (Sephadex G-25 resin, GE17-0851-01, Cytiva) were used to remove free biotin from samples and equilibrated according to the manufacturer’s instructions (Gravity Protocol) using 150 mM NaCl RIPA lysis buffer. 2 mL crude extract on column were eluted by 2.5 mL of 150 mM NaCl RIPA. Few microliters were collected to measure protein concentration of the extracts before and after desalting by performing Bradford assay (BioRad protein assay, Microassay procedure). Streptavidin Sepharose High Performance beads (GE Healthcare) were equilibrated twice with 1 mL 150 mM NaCl RIPA lysis buffer for 1 minute at RT. 70 µL of Dynabeads MyOne Streptavidine C1 (Invitrogen) were added to 5 mg of proteins with 1% protease inhibitor cocktail and incubated on a rotating wheel overnight at 4°C. Beads were then sequentially washed once with 1 mL of wash buffer 1 (2% SDS, 0.1% Triton), wash buffer 2 (50 mM HEPES pH 7.5, 500 mM NaCl, 1 mM EDTA, 0.5% sodium deoxycholate, 1% Triton), wash buffer 3 (10 Mm Tris-HCl pH 7.4, 250 mM LiCl, 1 mM EDTA, 0.5% sodium deoxycholate, 0.5% NP-40), and twice with wash buffer 4 (50 mM Tris-HCl pH 7.5, 50 mM NaCl, 0.1% NP-40).

To prepare samples for mass spectrometry analysis, beads were washed twice with 1 mL of 50 mM Tris-HCl pH 7.5, and transferred to a new tube and washed again with 1 mL Tris-HCl pH 7.5, 2 M urea. For the first tryptic digestion step, previous wash buffer is removed and 80 µL of Trypsin Buffer (50 mM Tris-HCl pH 7.5, 1 M urea, 1 mM DTT, 0.4 µg trypsin) is added to the beads and incubated at 25°C with agitation. After 3 h, beads were transferred to a new tube and washed twice with 60 µL of 50 mM Tris-Hcl ,1 M urea. Supernatants were collected to a new tube, DTT was added to a final concentration of 4 mM, and incubated at 25°C for 30 minutes with agitation. For alkylation of cysteine residues, 10 mM of iodoacetamide was added to the samples and incubated for 45 minutes at 25°C. For the final trypsin digestion step, 0.5 µg of trypsin was added to the samples and incubated overnight at 25°C. The following day, samples are acidified using 1% formic acid. Desalting of samples was carried out using OMIX C18 tips (10-100 µL, Agilent), previously activated and equilibrated twice using 200 µL Buffer B2 (0,1% formic acid, 50% acetonitrile) and four times using 200 µL Buffer A2 (0.1% formic acid). Samples were fixated with 8 suction-discharges and washed 10 times with 200 µL Buffer A2 before final elution with Buffer B2 in a new tube. Samples were completely dried using SpeedVac and then stored at -80°C. 4 independent replicates were treated and analyzed.

### TurboID nanoLC-MS/MS analyses

Biotinylated proteins were eluted from beads using biotin and digested with trypsin. Peptide samples were resuspended in 14 µL of 2% Acetonitrile, 0.05% TFA, vortexed and sonicated for 10 min before injection. Peptide mixtures were analysed by nano-LC-MS/MS using nanoRS UHPLC system (Dionex, Amsterdam, The Netherlands) coupled to a Q-Exactive Plus mass spectrometer (Thermo Fisher Scientific, Bremen, Germany). Five microliters of each sample were loaded on a C18 pre-column (5 mm × 300 *µ*m; Thermo Fisher) at 20 *µ*L/min in 5% acetonitrile, 0.05% trifluoroacetic acid. After 5 min of desalting, the pre-column was switched on line with the analytical C18 column (15 cm × 75 µm; Reprosil C18 in-house packed) equilibrated in 95% of solvant A (5% acetonitrile + 0.2% formic acid in water) and 5% of solvant B (80% acetonitrile + 0.2% formic acid in water). Peptides were eluted using a 5-50% gradient of B during 105 min at a 300 nL/min flow rate. The Q-Exactive Plus was operated in data-dependent acquisition mode with the Xcalibur software. Survey scan MS spectra were acquired in the Orbitrap on the 350-1500 *m/z* range with the resolution set to a value of 70 000. The ten most intense ions per survey scan were selected for HCD fragmentation, and the resulting fragments were analysed in the Orbitrap with the resolution set to a value of 17 500. Dynamic exclusion was used within 30s to prevent repetitive selection of the same peptide.

### Protein identification and quantification

Acquired MS and MS/MS data were searched with Mascot (version 2.8.0.1, http://matrixscience.com) against a custom-made database containing Turbo-ID proteins and TAIR database (release 20221110; 27,416 entries). The search included methionine oxidation, N-ter acetylation, S/T/Y phosphorylation as variable modifications and carbamidomethylation of cysteine as a fixed modification. Trypsin was chosen as the enzyme and 2 missed cleavages were allowed. The mass tolerance was set to 10 ppm for the precursor ion and to 20 mmu for fragment ions. Raw MS signal extraction of identified peptides was performed across different samples. Validation of identifications was performed through a false-discovery rate set to 1% at protein and peptide-sequence match level, determined by target-decoy search using the in-house-developed software Proline software version 2.1 (http://proline.profiproteomics.fr/) (Bouyssié et al., 2020). Raw MS signal extraction of identified peptides was performed with Proline across different samples. For all comparisons, statistical analysis was applied to the abundance values.

Proteins with a p-value from t-test < 0.05 and a ratio of average normalized area < 0.5 and > 2 were considered significant. Volcano plots were drawn to visualize significant protein abundance variations between two conditions, control and assay. They represent log10 (p-value) according to the log2 ratio. All the mass spectrometry data were deposited with the MassIVE repository with the dataset identifier: MSV000100915.

### Localization/colocalization confocal microscopy analyses

Plants were transferred from plate to liquid MS/2 medium with or without non-iron metal provision for 4 hours before observation. Plant samples were mounted in water and root tips were imaged on Leica SP8 confocal microscope using the 488 nm, 514 nm and 561 nm lasers to image GFP, mCit and mChe, respectively. Laser intensity settings were kept constant in individual sets of experiments to allow for comparison of expression and localization of reporter proteins. Quantification of total fluorescence intensity in roots, at the plasma membrane, in intracellular compartments and colocalization analyses were performed using ImageJ/Fiji (https://imagej.net/software/fiji/) as previously described (Dubeaux et al., 2018; Spielmann et al., 2023).

### TriFC protein:protein interaction assay

Emissions of TriFC-reconstituted mCit and the constitutively-expressed MyrPalm-mChe were collected in sequential mode. Mean fluorescence intensity was determined with ImageJ/Fiji (Measure tool) for both mCit and mChe channels following automatic selection of the plasma membrane region. This selection was based on the MyrPalm-mChe plasma membrane marker (pEAQ-HT-DEST1:MyrPalm-mCherry (Sainsbury et al., 2009)). mCit/mChe fluorescence ratios were then calculated, and statistical analyses were performed using GraphPad Prism as described in the legends to figures.

### Immunoprecipitation and western blot analyses

Immunoprecipitation (IP) experiments were carried out as previously described (Martín-Barranco et al., 2020). Approximately 1 g of fresh roots or *N.benthamiana leaves* were collected from 14-day-old seedlings or 48-hour-infiltrated *N.benthamiana*. Samples were ground in liquid nitrogen and resuspended in 3 volumes of cold solubilization buffer (50 mM Tris-HCl pH 7.5, 150 mM NaCl, 5mM EDTA, 1% [w/v] n-dodecyl-β-D-maltoside (DDM), 1% Plant-specific Protease Inhibitor Coktail (PIC)). Samples were centrifuged twice at 3,800 g for 10 minutes at 4°C, and placed on rotor wheel for solubilization of membrane proteins for at least 1 hour. Samples were then centrifuged 1 hour at 20, 000 g at 4°C for immunoprecipitation or 1 hour at 100,000 g at 4°C for coimmunoprecipitation. mCit-based immunoprecipitation was performed using µMACS GFP isolation kit (Miltenyi Biotec). RFP-based immunoprecipitation was performed using RFP-Trap_MA magnetic beads (Chromotek).

For protein detection, the following antibodies/probes were used: Monoclonal anti-GFP horseradish peroxidase-coupled (Miltenyi Biotech 130-091-833, 1/5,000), anti-ubiquitin P4D1 (Millipore 05-944, 1/2,500), anti-IRT1 (Agrisera AS11 1780, 1/5000), anti-FLAG M2 (Sigma-Aldrich, F1804, 1/2500), anti-RFP (Abcam AB34767, 1/5000), and avidin probe coupled to horse radish peroxidase (Sigma-Aldrich A3151, 1/5000).

### Endosome quantification in N. benthamiana

*N. benthamiana* plants were grown for 4 weeks in soil at 22°C with 16 hours light and 55% humidity and then infiltrated with *A.tumefaciens* GV3101 strain.

Treatment with the control solution (infiltration solution without metals, standard conditions) or non-iron metal solution (infiltration solution with -/+++ non-iron metal conditions) was done 48 hours after *A.tumefaciens* were infiltrated. Plants were analyzed by confocal microscopy after 3 hours of treatment using Leica SP8, with the 63X-oil-immersion objective, and images were collected at 1024x1024 pixel resolution. Endosome analysis was done using ComDet plugin from the Fiji software. ROIs were drawn around each cell and particles were detected with a maximum size of 4 pixels and an intensity threshold of 5.

### Statistical analyses

Statistical tests used, numbers of biological replicates, sample size in each biological replicate, and significance are described in the legend to the figures. All statistical tests were performed and graphs constructed using GraphPad Prism 8.

## Author contribution (CRediT)

L.P. : Methodology, Investigation ; S.F. Methodology, Investigation ; B.F. : Formal Analysis ; C.P. : Methodology, Investigation, Formal Analysis ; V.C. : Methodology, Supervision, Writing - Review & Editing ; G.V. : Conceptualization, Funding Acquisition, Project Administration, Supervision, Writing - Original Draft Preparation, Writing - Review & Editing. J.N. : Conceptualization, Methodology, Investigation, Formal Analysis, Validation, Supervision, Writing - Review & Editing.

## Acknowledgements

We would like to thank the imaging facility from FRAIB for their assistance.

## Data availability statement

All relevant data can be found within the manuscript and its supporting materials. Constructs and plant lines generated will be made available upon request to the lead contact.

