## Supplemental Figures and Tables for "TurboID-based proteomic profiling reveals proxitome of the IRT1 metal transporter and new insight into metal uptake regulation in plants"

Supplemental Figure 1. Impact of biotin addition on the non-iron metal-induced endocytosis and ubiquitination of IRT1.

(A) Confocal microscopy images of 7-day-old plants expressing IRT1::IRT1-mCit treated with different non-iron metal conditions, as established for TurboID, and subjected to mock or 200 µM biotin addition for 3 hours. Scale bar, 10 µm.

(B) IRT1 ubiquitination profile. Plants expressing UBI10:IRT1-mCit were grown as in (A) a,d subjected to immunoprecipitation using anti-GFP antibodies on solubilized root protein extracts. Immunoblotting was carried out with anti-GFP or anti-ubiquitin antibodies (P4D1). Inputs and immunoprecipitated fractions are shown. IB, immunoblotting; IP, immunoprecipitation. The stain free signal is used as loading control for inputs.

Alt Text : Microscopy data and western blot characterizing the impact of biotin on IRT1 localization and ubiquitination.

Supplemental Figure 2. Establishing non-iron metal conditions for TurboID.

(A) Confocal microscopy analysis of 7-day-old PIN2::IRT1-mCit plants treated with different non-iron metal conditions for 3 hours. Scale bar, 10 µm.

(B) Quantification of the plasma membrane to intracellular fluorescence ratio of plants coexpressing expressing PIN2::IRT1-mCit grown as in (A). Experiments were done in triplicates where five cells from three independent roots were imaged. Error bars represent standard deviation. Asterisks indicate significant (one-way ANOVA, Bonferroni's multiple comparisons test, ****P <0.0001 ; ns, not significant).

(C) Quantification of total fluorescence intensity of experiment in (A). Error bars represent standard deviation. Asterisks indicate significant (one-way ANOVA, Bonferroni's multiple comparisons test, ****P <0.0001 ; ns, not significant).

Alt Text : Microscopy data and graphs evaluating the experimental setup used for TurboID analyses.

Supplemental Figure 3. Evaluation of the interaction between IRT1 and NHX6.

(A) Principle of Trimolecular fluorescence complementation (TriFC). Interaction between IRT1-ALFA coupled to ALFA NB-mCitN and RGLG2-mCitC allows reconstitution of mCit fluorescence.

(B) Trimolecular fluorescence complementation (TriFC) assay in *Nicotiana benthamiana* leaves coexpressing ALFA NB-mCitN with NHX6-mCitC in presence (left) or absence (Ø, right) of IRT1-ALFA. The absence of IRT1-ALFA (Ø) serves as negative control. Representative images depicting the positive control interaction with AHA2 conducted simultaneously are shown in Figure 3.

(C) Ratiometric quantification of the mCitrine/MyrPalm-mCherry fluorescence signal ratios from confocal microscopy images of *N. benthamiana* shown in (B). Box-and-whisker plots indicate the median (line), interquartile range (box), 1.5 interquartile range (whiskers), and mean values ('+' symbol). Experiments were done in triplicates where six cells from two independent leaves were imaged. Data were analyzed by two-way ANOVA followed by Bonferroni multiple comparison post-test. Statistically significant differences between combinations (presence *vs* absence of IRT1-ALFA) are indicated by letters ($P<0.05$). Tested interactions were statistically different from negative controls. No statistical difference (n.s.) was observed among all tested interactors.

Alt Text : Illustration, microscopy data and graphs showing the ability of IRT1 to interact with NHX6 by TriFC.

Supplemental Figure 4. Impact of RGLG2 overexpression on IRT1 endocytosis.

(A) Representative confocal microscopy images of epidermal cells from *N.benthamiana* leaves transiently coexpressing UBI10::IRT1-mCit or its mutated counterpart UBI10::$IRT1_{2KR}$- mCit with UBI10::RGLG2-mChe or UBI10::$RGLG2_{2CS}$-mChe. Images were taken 3 hours after infiltration with control treatment (i.e. without non-iron metals, standard conditons) or after infiltration with non-iron metal excess treatment (non-iron metal treated). Maximum projection of 8-12 optical sections taken using 1 µm z-distance are shown. Scale bars, 10 µm.

(B) Quantification of intracellular particles per 100 $\mu m^2$ from plants shown in (A). Experiments were done in triplicates where three cells from three independent leaves were imaged. Asterisks indicate significant (two-way ANOVA , Sidakpost hoc test, ****$P < 0.0001$; ns, not significant).

Alt Text : Microscopy data and graphs quantifying the impact of RGLG2 overexpression on IRT1 localization and levels.

Supplemental Figure 5.

Venn diagram illustrating the overlap between IRT1 proximal proteins and proteins found in the IRT1 complex by AP-MS (Martín-Barranco et al., 2020).

Alt Text : Illustration of the overlap between the different proteomic-based approaches used to identify IRT1 partners.

Supplemental Table 1. Primers used for generating constructs

| Name | Forward primer sequence (5'-3') | Reverse primer sequence (5'-3') | Vector |
|---|---|---|---|
| attBTurboID-flag | GGGGACAAGTTTGTACAAAAAAGCAGGCTGA ACCATGAAAGACAATACTGTGCCTCTGAAG | GGGG AC CAC TTT GTA CAA GAA AGC TGG GTC TTTATCGTCATCGTCTTTGTAGTC | pDONR221 |
| TurboID-flag | GTTTAAACGCGGCCGCGGGAGGCGGTGGATCG AAAGAC | TCGACGCGGCCGCTTTATCGTCATCGTCTTTG | pDONR221-IRT1-insert1 |
| pUbi10 | GGGGACAACTTTGTATAGAAAA GTTGCTAGTCTAGCTCAACAGAGC | GGGGACTGCTTTTTTGTACAAACTT GCCTGTTAATCAGAAAAACT | pDONR-P4P1R |
| mCherry | GGGGACAACTTTGTATAATAAAGTTGCTT ACTTGTACAGCTCGTCCATGCCGCCGGTGGA | GGGGACAGCTTTCTTGTACAAAGTG GCTGTGAGCAAGGGCGAGGAG | pDPONR-P2RP3 |
| mCitrineN | GGGGACAGCTTTCTTGTACAAA GTGGCCATGGTGAGCAAGGGCGAG | GGGGACAACTTTGTATA ATAAAGTT GATTAGGCCATGATATAGACGTTGTGG | pDPONR-P2RP3 |
| mCitrineC | GGGGACAGCTTTCT TGTACAAA GTGGCCGACAAGCAGAAGAACGGCATC | GGGGACAACTTTGTATAATAAAGTT GATTACTTGTACAGCTCGTCCATG | pDPONR-P2RP3 |
| mCitrine | GGGGACAGCTTTCTTGTACAAA GTGGCTATGGTGAGCAAGGGCGAG | GGGGACAACTTTGTATAATAAAGTT GCTTACTTGTACAGCTCGTCCATGCCG | pDPONR-P2RP3 |
| pNHX5 | GGGGACAACTTTGTATAGAAAAGTT GCGAGATGTTCAATAAGCATCTCCA | GGGG AC TGC TTT TTT GTA CAA ACT TGCTCAGATTTGAGATTGGACCA | pDONR-P4P1R |
| NHX5 CDS | GGGGACAAGTTTGTACAAAAAAGCAGGC TGAACCATGGAGGAAGTGATGATTTCTC | GGGG AC CAC TTT GTA CAA GAA AGC TGG GTCCTCCCCATCTCCATCTCCAT | pDONR221 |
| pRGLG2 | GGGGACAACTTTGTATAGAAAAGTT GCCGATCCTCTGTGGCAAGGACTAG | GGGG AC TGC TTT TTT GTA CAA ACT TGC CAAACTTAACAAAAAACTAA | pDONR-P4P1R |
| RGLG2 CDS | GGGGACAAGTTTGTACAAAAAAGCAGGC TGAACC ATGGGGACAGGGAATTCTAA | GGGG AC CAC TTT GTA CAA GAA AGC TGG GTC GTAGAGCTTTATTCTTGTCTGGA | pDONR221 |
| NHX6 CDS | GGGGCGCGCCATGTCGTCGGAGCTGCAG | GGTTAATTAAGCCGCGGTTATTTAGATTTCC | pH7m34GW 2X35S:AscI-PacI:mCitC |
| RGLG2 C425,428S | ACCAGCTATCTCCGATTTCTTTGAGCA | TGCTCAAAGAAATCGGAGATAGCTGGT | Mutagenesis pDONR221-RGLG2 |
| RGLG2 C454,457S | TTCAAATGTCTCCGATTTCCCGTGCACCAATC | GATTGGTGCACGGGAAATCGGAGACATTTGAA | Mutagenesis pDONR221-RGLG2 |
| LTI6B CDS | GGGGACAGCTTTCTTGTACAAAGTGGCCATGA GTACAGCCACTTTCG | GGGGACAACTTTGTATAATAAAGTTGATTACTTGGT GATGATATAAAGAG | pDPONR-P2RP3 |

Supplemental Table 2. List of constructs generated or used

| Name | Source |
|---|---|
| *pDONR221-IRT1-notI position1* | Neveu et al., 2025 |
| *pDONR221-IRT1 TurboID position1* | This study |
| *pDONR221-NHX5* | This study |
| *pDONRP2RP3-mCherry* | Marquès-Bueno et al., 2016 |
| *pDONR221-RGLG2* | This study |
| *pK7m34GW UBI:IRT1-mCit* | Spielmann et al., 2022 |
| *pG7m34GW PIN2:IRT1-mCit* | Dubeaux et al., 2018 |
| *pG7m34GW IRT1:ALFAtag - IRT1 position 1* | Neveu et al., 2025 |
| *pH7m34GW 2X35S:AscI-FRO2-PacI:mCitC* | Neveu et al., 2025 |
| *pH7m34GW 2X35S:AscI-AHA2-PacI:mCitC* | Neveu et al., 2025 |
| *pB7m34GW 2X35S:Alfa-nanobody:mCitN* | Neveu et al., 2025 |
| *pDONRP2RP3-Lti6b w STP* | Neveu et al., 2025 |
| *pDONRR221 citrine* | Marquès-Bueno et al., 2016 |
| *PDONRP4P1R-UBI10* | Marquès-Bueno et al., 2016 |
| *pDONR221-NHX6* | This study |
| *pDONR221-TurboID* | This study |
| *pDONRP4P1R-NHX5* | This study |
| *pDONRP4P1R-RGLG2* | This study |

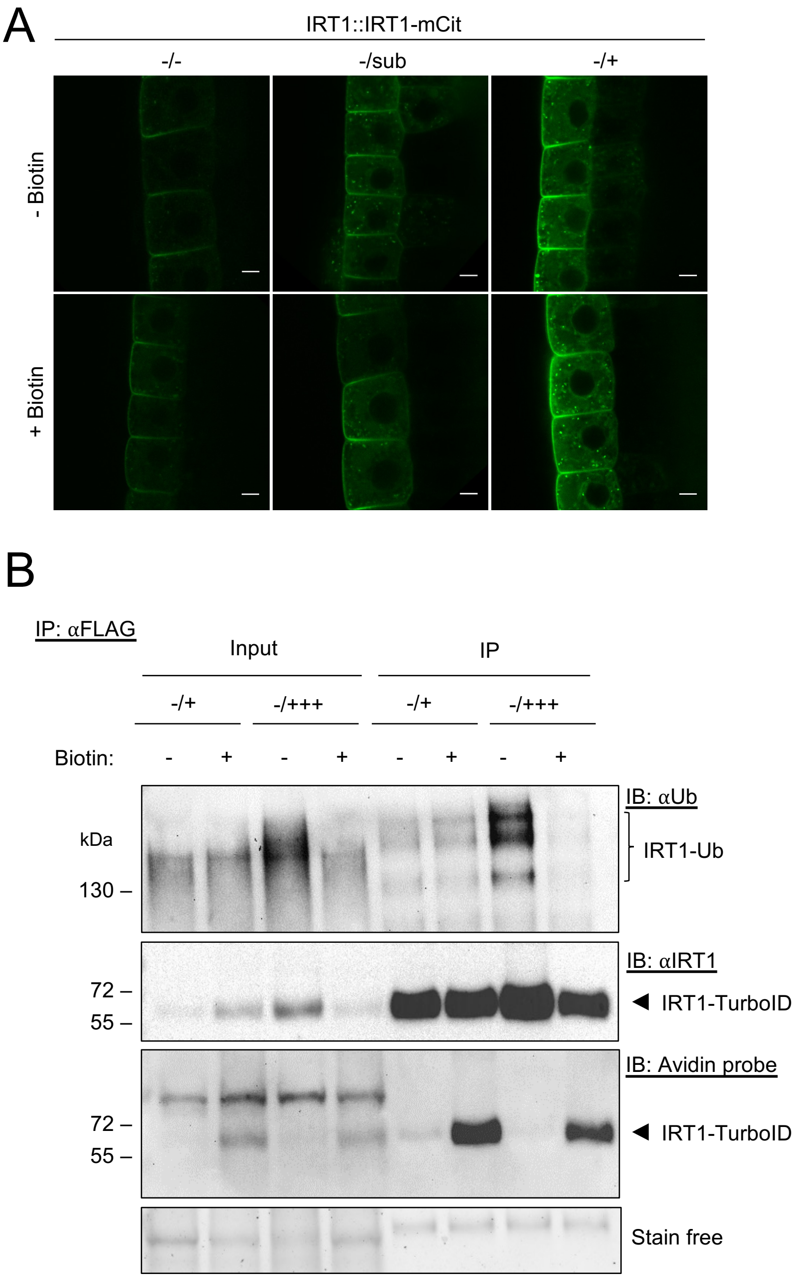
A
IRT1::IRT1-mCit
-/-
-/sub
-/+
- Biotin
+ Biotin
B
IP: αFLAG
Input
IP
-/+
-/+++
-/+
-/+++
Biotin:
-
+
-
+
-
+
-
+
kDa
130 –
IB: αUb
IRT1-Ub
72 –
55 –
IB: αIRT1
IRT1-TurboID
72 –
55 –
IB: Avidin probe
IRT1-TurboID
Stain free

Figure S1

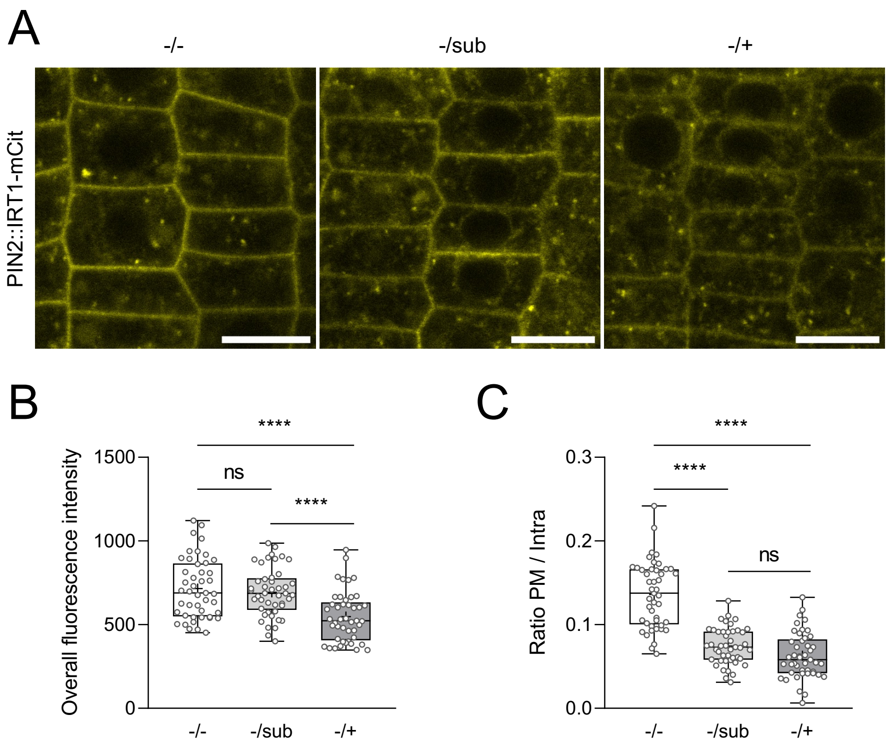
A
-/-
-/sub
-/+
PIN2::IRT1-mCit
B
Overall fluorescence intensity
1500
1000
500
0
ns
****
****
-/-
-/sub
-/+
C
Ratio PM / Intra
0.3
0.2
0.1
0.0
****
****
ns
-/-
-/sub
-/+

Figure S2

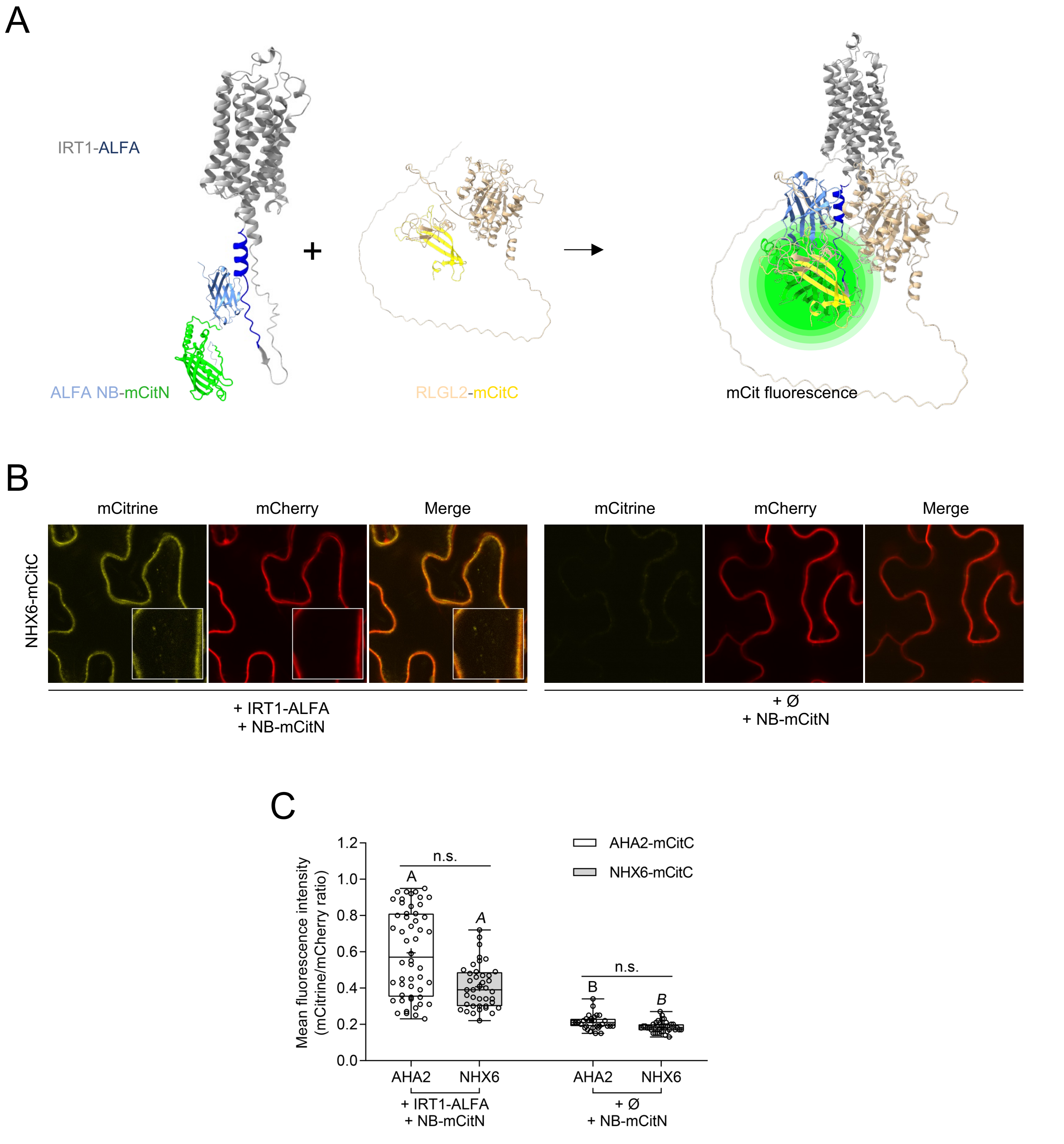
A
IRT1-ALFA
ALFA NB-mCitN
RLGL2-mCitC
mCit fluorescence
B
mCitrine
mCherry
Merge
NHX6-mCitC
+ IRT1-ALFA
+ NB-mCitN
+ Ø
+ NB-mCitN
C
Mean fluorescence intensity (mCitrine/mCherry ratio)
AHA2-mCitC
NHX6-mCitC
n.s.
A
B
AHA2
NHX6
+ IRT1-ALFA
+ NB-mCitN
+ Ø
+ NB-mCitN

Figure S3

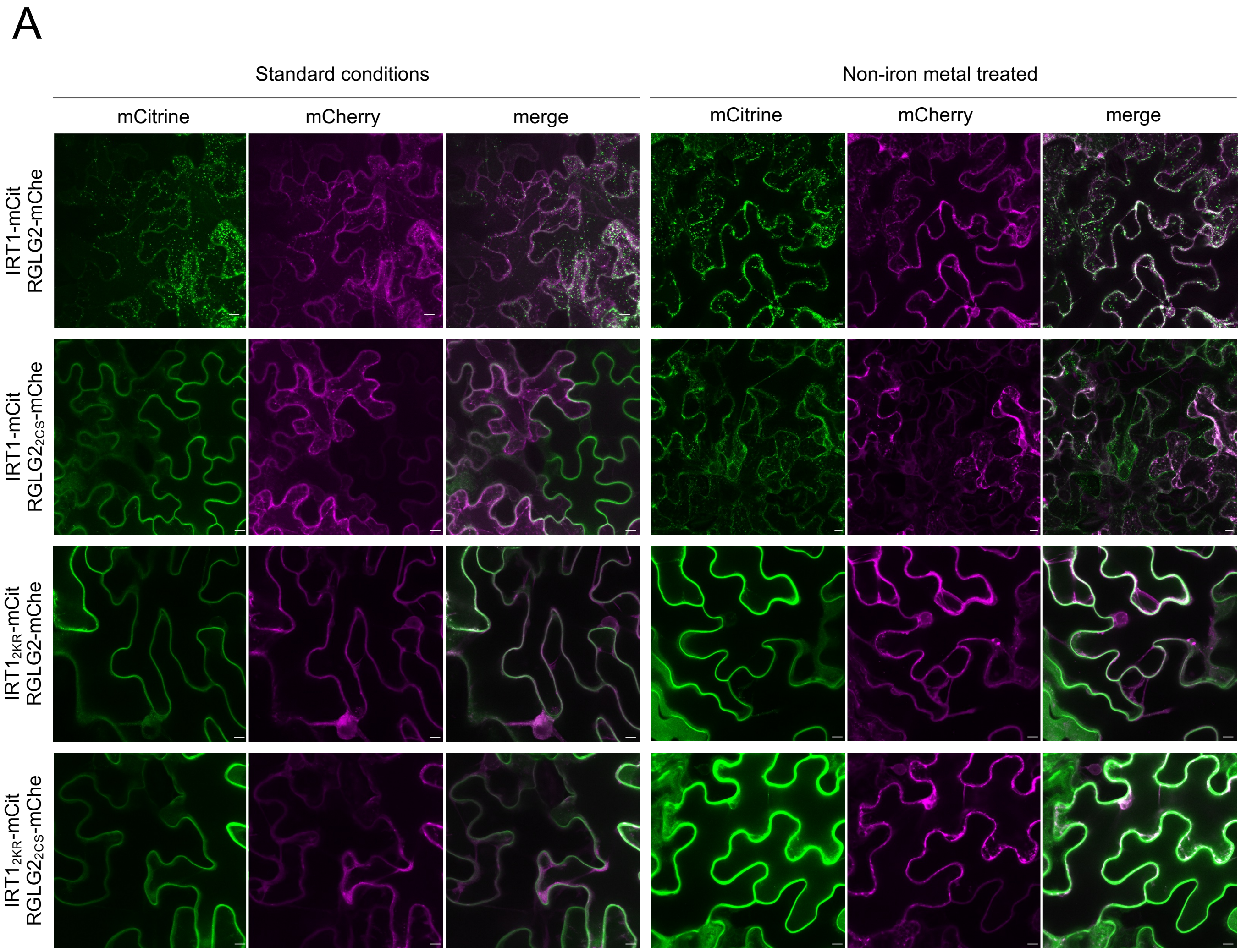
A
Standard conditions
Non-iron metal treated
mCitrine
mCherry
merge
mCitrine
mCherry
merge
IRT1-mCit
RGLG2-mChe
IRT1-mCit
RGLG2_2CS-mChe
IRT1_2KR-mCit
RGLG2-mChe
IRT1_2KR-mCit
RGLG2_2CS-mChe

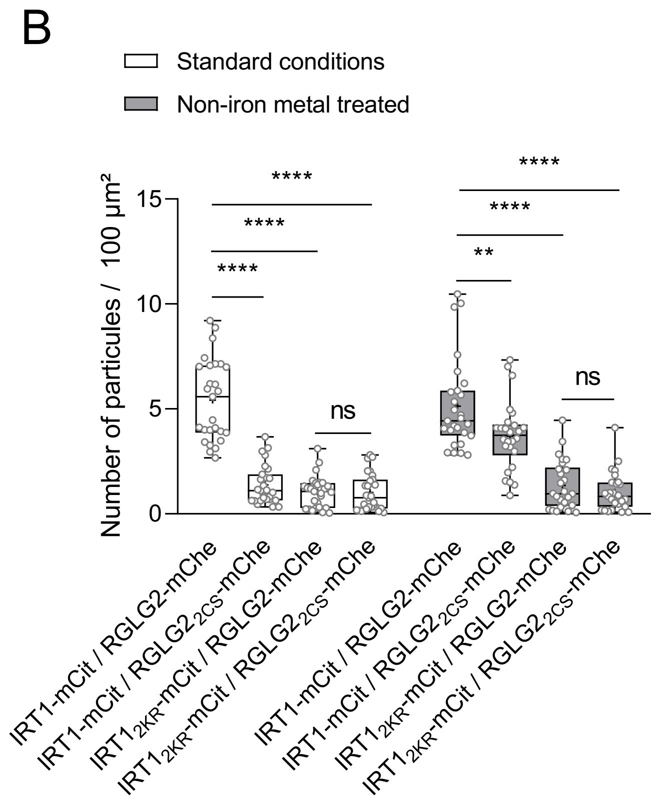
B
Standard conditions
Non-iron metal treated
Number of particules / 100 µm²
15
10
5
0
****
****
****
ns
****
****
**
ns
IRT1-mCit / RGLG2-mChe
IRT1-mCit / RGLG2_2CS-mChe
IRT1_2KR-mCit / RGLG2-mChe
IRT1_2KR-mCit / RGLG2_2CS-mChe
IRT1-mCit / RGLG2-mChe
IRT1-mCit / RGLG2_2CS-mChe
IRT1_2KR-mCit / RGLG2-mChe
IRT1_2KR-mCit / RGLG2_2CS-mChe

Figure S4

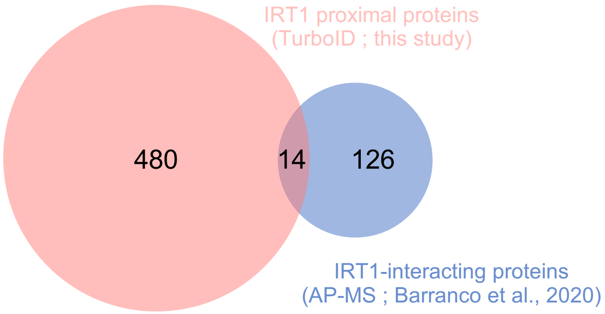
IRT1 proximal proteins
(TurboID ; this study)
480
14
126
IRT1-interacting proteins
(AP-MS ; Barranco et al., 2020)

Figure S5
